# Investigating the Effect of Flow Compensation and Quantitative Susceptibility Mapping Method on the Accuracy of Venous Susceptibility Measurement

**DOI:** 10.1101/2021.04.14.439812

**Authors:** Ronja C. Berg, Christine Preibisch, David L. Thomas, Karin Shmueli, Emma Biondetti

## Abstract

Quantitative susceptibility mapping (QSM) is a promising non-invasive method for obtaining information relating to the oxygen metabolism. However, the optimal acquisition sequence and QSM reconstruction method for reliable venous susceptibility measurements are unknown. Full flow compensation is generally recommended to correct for the influence of venous blood flow, although the effect of flow compensation on the accuracy of venous susceptibility values has not been systematically evaluated. In this study, we investigated the effect of different acquisition sequences, including different flow compensation schemes, and different QSM reconstruction methods on venous susceptibilities.

Ten healthy subjects were scanned with five or six distinct QSM sequence designs implementing different flow compensation schemes. All data sets were processed using six different QSM pipelines and venous blood susceptibility was evaluated in whole-brain segmentations of the venous vasculature and single veins. The quality of vein segmentations and the accuracy of venous susceptibility values were analyzed and compared between all combinations of sequences and QSM methods.

The influence of the QSM method on average venous susceptibility values was found to be 2.7 - 11.6 times greater than the influence of the acquisition sequence, including flow compensation. The majority of the investigated QSM reconstruction methods tended to underestimate venous susceptibility values in the vein segmentations that were obtained.

Using multi-echo gradient-echo acquisitions with monopolar readout gradients, we found that sequences without full flow compensation yielded venous susceptibility values comparable to sequences with full flow compensation. However, the QSM method had a great influence on susceptibility values and thus needs to be considered carefully for accurate venous QSM.

## 1 Introduction

Venous quantitative susceptibility mapping (QSM) aims to quantify the magnetic susceptibility (χ) of venous blood based on the phase of the gradient-recalled echo (GRE) magnetic resonance imaging (MRI) signal, which reflects the paramagnetic susceptibility of deoxygenated hemoglobin and venous oxygen saturation (SvO_2_) (Jain et al., 2012; Spees et al., 2001; Weisskoff and Kiihne, 1992). SvO_2_ is linearly related to χ of blood and can be calculated as (Weisskoff and Kiihne, 1992):

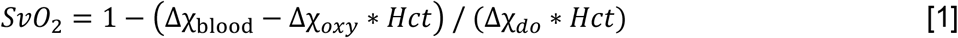

where Δχ_blood_ is the measured susceptibility difference between blood and water (or soft tissue), Δχ_*oxy*_ the constant susceptibility shift of fully oxygenated blood relative to water, Δχ_*do*_ the known susceptibility difference between fully oxygenated and fully deoxygenated red blood cells, and *Hct* the hematocrit.

Based on Equation [1], previous studies have successfully quantified SvO_2_ in healthy subjects (Fan et al., 2014; Ward et al., 2017; Xu et al., 2014) and patients with cerebrovascular disease (Biondetti et al., 2019; Fan et al., 2020; Schneider et al., 2020). For assessing brain oxygenation, QSM represents a non-invasive and more feasible alternative to the gold standard ^15^O PET (Mintun et al., 1984). Indeed, PET employs short-lived ^15^O radiotracers that require on-site production by a cyclotron, whereas QSM exploits the oxygenation status of hemoglobin, a naturally occurring endogenous contrast mechanism (Cho et al., 2020; Fan et al., 2015; Kudo et al., 2016). However, for venous QSM, neither the acquisition sequence nor the image processing pipeline has yet been standardized (Bilgic et al., 2020; Langkammer et al., 2018).

Because of the known effects of moving spins on signal phase, it has been suggested that GRE images acquired for venous QSM require full first-order (i.e., velocity) flow compensation (Brown et al., 2014). Indeed, in the absence of flow compensation, vessels containing flowing blood can appear displaced on GRE magnitude (Deistung et al., 2009), GRE phase (Xu et al., 2014), SWI (Deistung et al., 2009), and QSM (Biondetti et al., 2020) images, potentially affecting the accuracy of both image-based vessel segmentation and venous susceptibility estimation inside segmented vessels. Fully flow-compensated sequences aim to suppress the additional phase induced by flowing spins (velocities in veins: 10-25 cm/s) at each echo time (TE) and along all three signal encoding directions of the Cartesian k-space trajectory.

Acquiring multi-echo GRE images for QSM is desirable because it enables optimization of the signal-to-noise ratio (SNR) in multiple tissue types simultaneously (Haacke et al., 2015; Wu et al., 2012a) and fitting over echo times for accurate field map estimation for QSM (Biondetti et al., 2020). However, commercially available scanner options enable full flow compensation only for single-echo protocols or at the first TE of multi-echo protocols. Moreover, the longer TE images of a multi-echo GRE protocol are usually only flow-compensated along the frequency-encoding direction (Denk and Rauscher, 2010; Gilbert et al., 2012). Therefore, pulse sequence programming is required to enable acquisition of fully flow-compensated multi-echo GRE images (Deistung et al., 2009; Xu et al., 2014), limiting the use of these sequences in clinical settings.

In addition to image acquisition, the QSM reconstruction method is expected to impact on measurement accuracy for venous susceptibility. While optimization and standardization of QSM reconstruction algorithms and processing pipelines is an active area of research within the QSM community (Bilgic et al., 2020; Langkammer et al., 2018; Robinson et al., 2017; Schweser et al., 2017), only a few studies have proposed methods specifically optimized for venous QSM reconstruction (Biondetti et al., 2019; Haacke et al., 2015; Liu et al., 2013; Wei et al., 2015; Xu et al., 2014). Critically, the effect of distinct QSM reconstruction pipelines on the accuracy of venous QSM has never been systematically compared.

To systematically assess the effect of flow compensation as well as different QSM reconstruction methods on the accuracy of measured venous susceptibility values, we aimed to systematically compare different acquisition sequences as well as different QSM reconstruction pipelines. To this end, we set up distinct multi-echo GRE sequences with flow compensation applied at all TEs, at the first TE only, or without any flow compensation. Furthermore, because time constraints are known to limit the clinical applicability of multi-echo QSM sequences, we investigated the possibility of using state-of-the-art acceleration techniques, namely Compressed SENSE (Geerts-Ossevoort et al., 2018), compared to the standard SENSE technique (Pruessmann et al., 1999). To independently assess venous anatomy and facilitate vein segmentation, we also acquired phase contrast angiography (PCA) with velocity encoding targeting venous flow. With respect to QSM reconstruction pipelines, we focused primarily on methods featured in the most popular toolboxes available for QSM, since these can be employed by non-specialist users and are therefore feasible in clinical settings. To evaluate their performance, these non-optimized pipelines were compared to a QSM processing method that had been specifically optimized for venous QSM in a previous study (Biondetti et al., 2019) and to an iterative version of the same method (Karsa et al., 2020).

### 2. Methods

### 2.1. Subjects

This study was approved by the local medical ethical committee at the Klinikum rechts der Isar, Technical University of Munich (TUM). After providing informed written consent for participation in this study, ten healthy volunteers (five females, age range: 22-50 years, average age: 29 years) underwent MRI at the Department of Neuroradiology, Klinikum rechts der Isar, TUM.

To investigate reproducibility across different MRI systems from the same vendor, data from five healthy volunteers (3 females, age range: 30-54 years, average age: 44 years) that had been scanned previously in a pilot study were included in the analysis. The pilot study was performed at the Queen Square Multiple Sclerosis Centre (University College London (UCL), UCL Institute of Neurology, London, UK). This study was approved by the local ethics review board and all subjects provided written informed consent.

### 2.2. Image acquisition

Data acquisition was performed on a Philips Ingenia Elition X 3 T MR system (Philips Healthcare, R5.6.1.0, Best, NL) using a 32-channel head coil. All imaging sequences for QSM and PCA were based on a 3D multi-echo GRE sequence. Images were acquired in transverse orientation with 1-mm isotropic resolution, 246 x 188 x 144 mm^3^ field-of-view (read x phase encoding (PE)_1_ x PE_2_), with frequency encoding in the anterior-posterior direction and a right-left primary phase encoding (fold-over) direction. SENSE (2-fold acceleration in the PE_1_ direction) was used as the standard imaging acceleration technique. All GRE sequences for QSM had monopolar (flyback) readout gradients enabled, flip angle 17°, and the shortest achievable repetition time (which was 43 or 44 ms for all sequences). Other acquisition parameters, namely the number of echoes and the readout bandwidth, were optimized in line with each sequence’s flow compensation scheme (see Table 1 for a detailed overview). Additionally, in one sequence, Compressed SENSE (CS) (Geerts-Ossevoort et al., 2018) was applied (4-fold acceleration) to evaluate the effect of reducing acquisition time by sparse sampling of k-space. The following flow compensation schemes were tested: full flow compensation for all echoes and along all encoding directions (Full-FC) as in a previous study (Xu et al., 2014); a conventional flow compensation scheme for the first echo only as implemented by the vendor using SENSE (TE1-FC) or Compressed SENSE (TE1-FC-CS); and no flow compensation with TEs matched either to the Full-FC sequence (No-FC) or with the maximum number of TEs within the TR (No-FC-7ech). Finally, to investigate the effect on multi-echo QSM accuracy of maximizing the SNR over the same TR, a sequence was implemented with no flow compensation and three TEs (No-FC-3ech), which is the minimum number of TEs for multi-echo QSM using the Morphology Enabled Dipole Inversion (MEDI) toolbox (see section 2.3.2). All sequences were run on all subjects except No-FC-3ech, which, because of time restrictions, was only tested on four subjects. Additionally, PCA data were acquired to independently locate and segment major cerebral veins. The GRE sequence implementation with full flow compensation was enabled by research software made available by the scanner manufacturer.

**Table 1:**
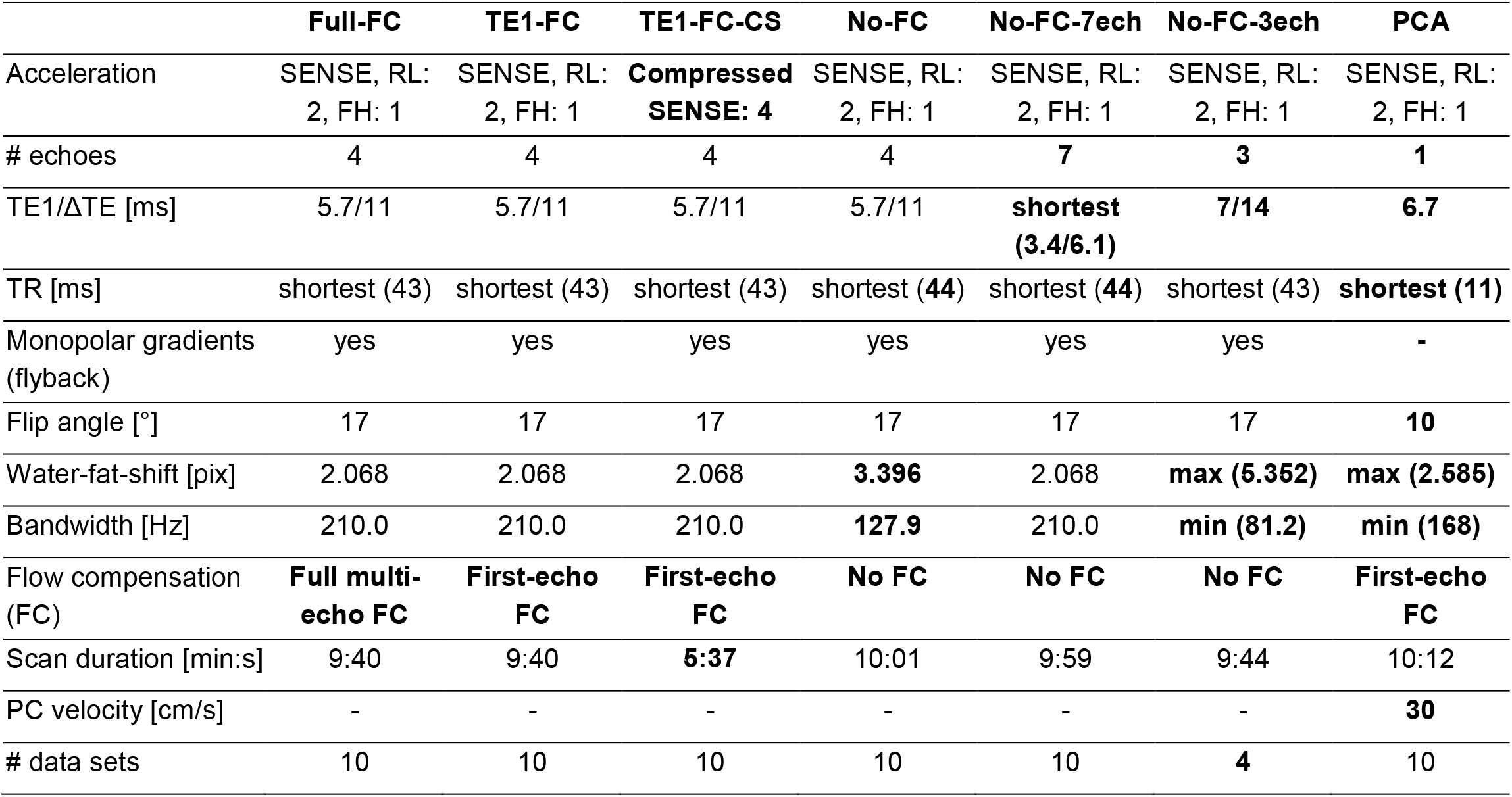
Key acquisition parameters and flow compensation schemes for all of the 3D GRE sequences for QSM and the 3D GRE phase contrast angiography (PCA). Parameters deviating from the most common value are marked in bold. Differences between sequences relevant for QSM include differences in the flow compensation scheme (influencing the bandwidth / water-fat-shift), the number of echoes and echo times, and the image acceleration technique (influencing the scan duration). Common parameters were 1 x 1 x 1 mm^3^ resolution, 246 x 188 x 144 mm^3^ field-of-view, transverse slice orientation, and right-left fold-over direction. TE: echo time, ΔTE: echo spacing, TR: repetition time, PC: phase contrast, RL: right-left, FH: foot-head.

The data of the pilot study had been acquired on a Philips Achieva 3 T system (software version R3.2.1) using a 32-channel head coil. The acquisition protocol comprised three of the sequences used in the main study: Full-FC, No-FC, and No-FC-7ech with identical parameter settings except for a 1.1 x 1.1 x 1.1 mm^3^ resolution and a 246 x 180 x 144 mm^3^ field-of-view.

### 2.3. Image Processing and Analysis

Where not stated otherwise, all data processing and analyses were performed using MATLAB (R2020a, The Mathworks, Natick, MA, United States).

#### 2.3.1. Brain mask calculation

For each subject, brain masks for QSM were calculated based on the longest-TE magnitude image from each sequence. This enabled exclusion of anatomical areas near air-tissue interfaces, which cause artifacts due to signal dropout at longer TEs. The magnitude images were segmented into tissue compartments using the “segment” module of the Statistical Parametric Mapping toolbox (SPM12; https://www.fil.ion.ucl.ac.uk/spm/software/spm12/) for MATLAB. Each brain mask was calculated as the sum of the gray matter, white matter, and cerebrospinal fluid tissue compartments. In each transverse slice, residual holes in the brain mask were filled using an in-house MATLAB function that evaluates each voxel in the mask in relation to its four direct neighbors. The superior sagittal sinus was added to the brain mask using ITK-SNAP’s semi-automated segmentation tool (Yushkevich et al., 2006) (www.itksnap.org) on the first-echo magnitude image as detailed in a previous study (Biondetti et al., 2019).

#### 2.3.2. QSM processing pipelines

Each 3D QSM data set was processed using six different QSM reconstruction pipelines (see Figure 1 for a schematic overview). Three methods were based on two publicly available toolboxes: the MEDI toolbox (version 11/2017; http://pre.weill.cornell.edu/mri/pages/qsm.html) and the Susceptibility Tensor Imaging (STI) Suite (v3.0_05/2017; https://people.eecs.berkeley.edu/~chunlei.liu/software.html), both with default settings applied. Additionally, we tested total generalized variation (TGV) (Langkammer et al., 2015), a one-step algorithm, and two Tikhonov (TIK)-based QSM reconstruction methods, the latter having been optimized and applied previously for venous QSM (Biondetti et al., 2020; Biondetti et al., 2019). All methods except TGV used the same brain mask, calculated as described above. Calculated susceptibility values were not referenced to a specific brain tissue to avoid combining the effects of acquisition or QSM method in a reference ROI with those in venous ROIs.

**Figure 1:**
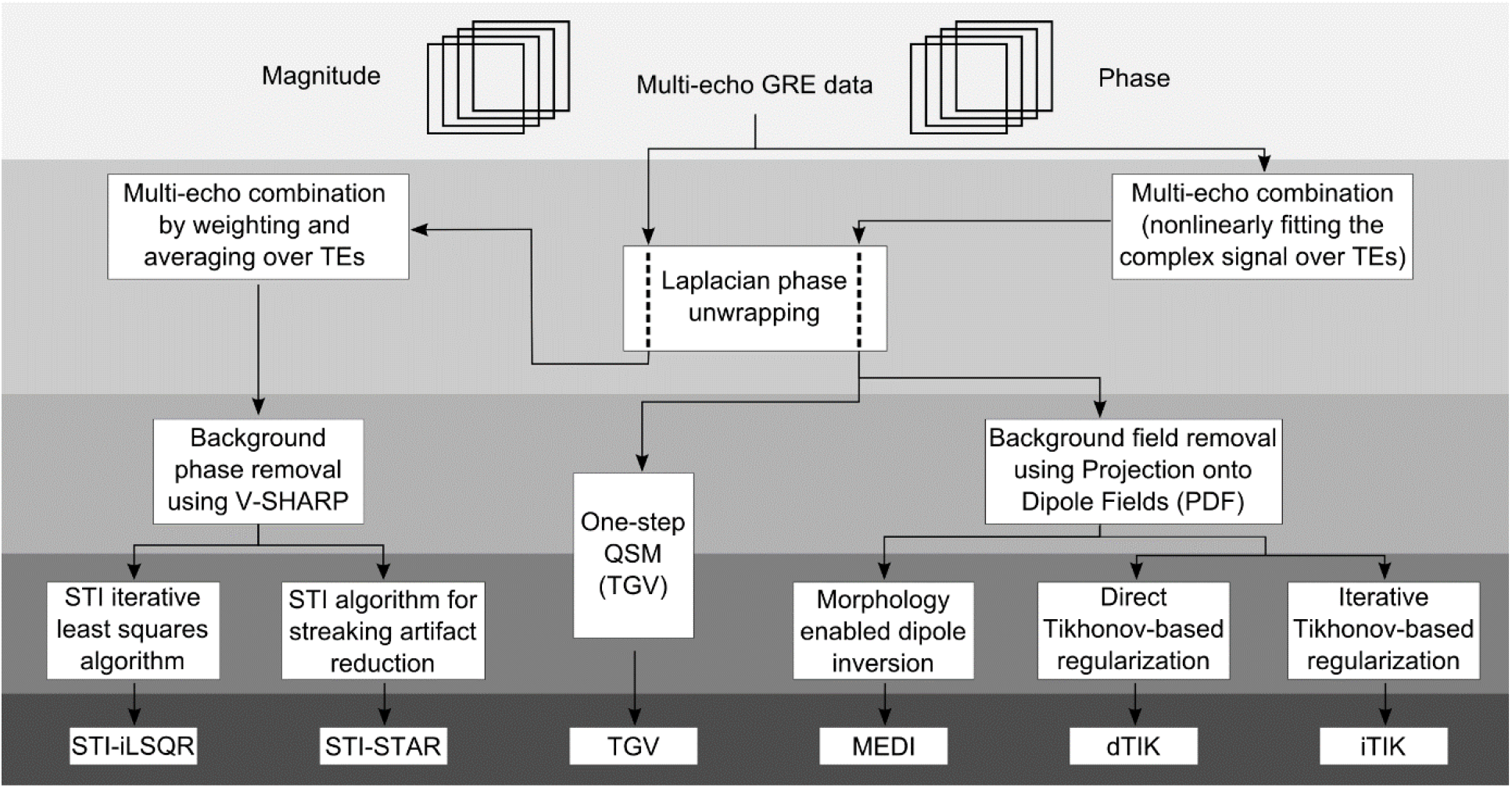
Schematic diagram of the processing steps in each QSM reconstruction pipeline. The input data consisted of magnitude and phase images from each of the different gradient-echo sequences listed in Table 1. Multi-echo combination was performed after Laplacian phase unwrapping when using STI Suite and before phase unwrapping in all other pipelines. In TGV, background field removal and field-to-susceptibility inversion were performed within the same step. All other QSM pipelines performed these two steps separately.

For multi-step processing using the MEDI toolbox, the total field map was calculated by fitting the complex GRE signal over TEs. Phase unwrapping was performed using a Laplacian-based technique (Schofield and Zhu, 2003), and background fields were removed via projection onto dipole fields (de Rochefort et al., 2010; Liu et al., 2011). Local field-to-susceptibility inversion was performed using the Morphology Enabled Dipole Inversion “MEDI_L1” (Liu et al., 2012; Liu et al., 2011) method with a reduced radius of 3 for the spherical mean value operator “SMV” to avoid excessive erosion of the brain mask, and otherwise default values. This included a default value of 1000 for the regularization parameter (recommended by the developers for brain applications and verified by checking several values on these data), since the toolbox did not provide any options for optimizing this parameter.

In the STI Suite, Laplacian-based phase unwrapping was applied and the multi-echo unwrapped phase images were then combined via TE-weighted averaging (Li et al., 2015; Wei et al., 2015). Background fields were removed using sophisticated harmonic artifact reduction for phase data with variable kernel sizes (V-SHARP) (Li et al., 2011; Wu et al., 2012b). Local field-to-susceptibility inversion was performed using two distinct methods: iterative least-squares (iLSQR) (Li et al., 2011) and an algorithm specifically designed for reducing streaking artifacts potentially arising from high-susceptibility sources such as large veins (streaking artifact reduction, STAR) (Wei et al., 2015).

One-step TGV-based QSM (Langkammer et al., 2015) was performed using Singularity (Sylabs Inc., Albany, CA, USA; https://github.com/CAIsr/qsm) and the default parameter values *α*_1_, *α*_0_ = 0.0015, 0.0005 based on the criterion that *α*_1_: *α*_0_ = 3:1 is optimal for medical imaging applications (Knoll et al., 2011). The inputs for TGV QSM were the field map combined using the nonlinear fitting function in the MEDI toolbox (Liu et al., 2013) and the echo spacing.

The input for the Tikhonov-based QSM calculations was the local field map calculated by the MEDI-based pipeline. Local field-to-susceptibility inversion was performed using direct Tikhonov regularization (dTIK; https://xip.uclb.com/i/software/mri_qsm_tkd.html) as well as an iterative implementation (iterative Tikhonov, iTIK; https://xip.uclb.com/i/software/mri_qsm_tkd.html), both with correction for susceptibility underestimation. For dTIK, the average optimal regularization parameter “alpha” (*α*_*dT*_) across subjects was calculated separately for each acquisition sequence using the L-curve method (Hansen and O’Leary, 1993). The values for *α*_*dT*_ ranged between 0.070 and 0.074 and can be found in Table S1 in the supplementary material. For iTIK, an empirical fixed value of *α*_*iT*_ = 0.05, recommended by the developer for brain applications, was used as the regularization parameter.

For each acquisition sequence, a minimum-size brain mask was calculated as the intersection of the masks calculated by all of the distinct QSM pipelines.

#### 2.3.3. Whole-brain vessel filtering

The automated multiscale vessel filtering (MVF) method (Bazin et al., 2015) (version 3.0.7) from the JIST-LayoutTool (v1.8, 08/2013; https://www.nitrc.org/projects/jist/) of MIPAV (v8.0.2, 02/2018; https://mipav.cit.nih.gov/) was used for whole-volume vessel segmentation of the susceptibility maps from each QSM pipeline. The segmentation was performed using default parameter settings including a recursive ridge filter and a scale number (kernel size) equal to 4 (Biondetti et al., 2019).

#### 2.3.4. Single-vein segmentation

Based on the PCA venogram, semi-automated segmentation of major representative veins (superior sagittal sinus (SSS), straight sinus (StrS), transverse sinuses (TraS), and internal cerebral veins (ICVs)) was performed using ITK-SNAP’s “active contour segmentation mode” (Bettoni et al., 2018; Law and Chung, 2012). Here, the TraS segment comprised the transverse sinuses, extending into the sigmoid sinuses and the internal jugular veins. It was defined to investigate the global performance of venous QSM in the inferior part of the brain. The PCA-based venous regions of interest (ROIs) were aligned with each susceptibility map using SPM12 via rigid alignment of the PCA magnitude image with the corresponding first-echo GRE magnitude image. To mitigate partial volume effects between veins and surrounding brain tissue, the co-registered ROIs of larger veins (i.e., SSS, StrS, and TraS) were eroded with a spherical shaped structural element by one voxel for further analyses.

#### 2.3.5. Quantitative analysis

All vein segmentations (whole-brain and single-veins) were multiplied with the minimum-size brain mask. The average and standard deviation of susceptibility values were calculated in both the masked whole-brain MVF-based and single-vein segmentations. For each susceptibility map, venous density was calculated as the fraction of MVF-segmented venous voxels over the total number of voxels in the brain mask.

To assess the accuracy of MVF-based automated segmentation, the non-eroded PCA-based single-vein ROIs were considered as a gold-standard reference and the fraction of venous voxels correctly detected within these ROIs by the MVF segmentation was calculated. Furthermore, the number of voxels that were “uniquely” segmented in only one of the six susceptibility maps was evaluated by comparing whole-brain segmentations of the susceptibility maps from each of the different QSM reconstruction pipelines for each sequence and subject. Similarly, the number of commonly segmented voxels was determined for each acquisition sequence as the number of voxels segmented in all six susceptibility maps reconstructed from that sequence. For each sequence and QSM method, the number of uniquely segmented voxels was then divided by the number of voxels commonly segmented in all six susceptibility maps from different QSM reconstruction methods for that sequence to obtain their ratio. Additionally, to assess the similarity between segmentation outcomes across susceptibility maps from different acquisition sequences and reconstruction methods, Sørensen-Dice similarity coefficients were calculated between automated whole-brain segmentations from different susceptibility maps. The coefficients were computed as (Dice, 1945) 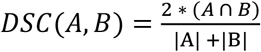,where |A| is the number of voxels in the segmentation of susceptibility map (A), |B| the number of voxels in the segmentation susceptibility map (B), and (*A* ∩ *B*) the number of voxels in the intersection of both segmentations. Finally, to analyze the similarity of the two segmentation methods, the intersection of the PCA-based segmentation of ICVs and the MVF-segmentation was used. To this end, the “bwconncomp” MATLAB function was utilized to find the largest connected component within the intersection map. The number of voxels within this largest connected component was then divided by the total number of voxels in the PCA-based ICVs segmentations.

To estimate the respective influence of QSM reconstruction methods and imaging sequences, venous susceptibilities from data acquired with different sequences but reconstructed with the same QSM reconstruction method were averaged to obtain a method-mean venous susceptibility value, 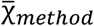. The maximal difference between the six method-mean venous susceptibility values 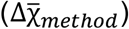 and the largest difference between the six sequence-mean venous susceptibility values 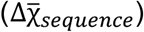 were determined, and their ratio was calculated to assess the relative importance of the two factors.

Venous oxygenation values were calculated according to Equation [1] with *Hct* = 0.4, Δχ_*do*_ = 0.27 * 4π ppm (Spees et al., 2001), and Δχ_*do*_ = -0.03 * 4π ppm (Weisskoff and Kiihne, 1992). 2.3.6.

#### 2.3.6. Statistical analysis

Statistical analyses were performed using SPSS (IBM Corp. Released 2019. IBM SPSS Statistics for Windows, Version 26.0. Armonk, NY). For whole-brain MVF-based and semi-automated single-vein segmentations, separate two-way repeated measures analyses of variance (ANOVAs) were applied to test whether there is a statistically significant difference in average venous susceptibility when using different imaging sequences and applying distinct QSM reconstruction pipelines. The Shapiro-Wilk test and Mauchly’s Test of Sphericity were applied to test and verify the normality of each data set’s distribution and sphericity, respectively, as requirements for performing the ANOVA. Effect sizes (η^2^) were calculated as (Lakens, 2013) 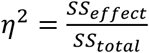 where *SS*_*effect*_ is the sum of squares of an effect (in this study, e.g., QSM method or sequence) and *SS*_*total*_ the total sum of squares (variability of venous susceptibility values in the study). Simple main effects were analyzed for all average venous susceptibilities of segmentations that showed a significant interaction between both effects (QSM method and sequence). Therefore, paired samples t-tests were run for all possible combinations of the six QSM reconstruction methods (15 per sequence) for the five sequences that were acquired in all ten subjects (75 combinations) and for all pairs of sequences per reconstruction method (60 combinations). The Benjamini-Hochberg procedure was then applied to control for the false discovery rate.

### 2.4. Data availability statement

In line with local ethics guidelines and subject privacy policies, the acquired data are only available via a request to the authors. Institutional policies require a formal data sharing agreement. The full MATLAB code applied for susceptibility map calculation is available upon request but all of the QSM reconstruction methods (Figure 1) are publicly available via the links above (section 2.3.2). Sharing of applied sequence modifications is limited by a nondisclosure agreement with the scanner manufacturer.

## 3. Results

### 3.1. Overall vessel appearance on QSM

Multi-echo GRE images acquired using the different acquisition sequences appeared similar on visual inspection (Figure 2). Likewise, differences between reconstructed susceptibility maps acquired using different flow compensation schemes appeared small on simple visual inspection (Figure 3). Instead, susceptibility maps reconstructed with the six different QSM reconstruction methods exhibited clear differences in overall tissue contrast, extent of brain erosion, also affecting SSS visibility, and vein delineation performance (Figure 3). Some streaking artifacts were found in the MEDI reconstructions (Figures 3, MEDI, sagittal slices), while the Tikhonov-based reconstructions seemed to contain residual low-frequency (background) field contributions (Figures 3, dTIK and iTIK). Generally, in the MEDI, TGV, and dTIK susceptibility maps, veins had a higher contrast relative to the surrounding brain tissue, but substantially decreased susceptibility values in the immediate vicinity of venous areas (Figure 3, arrowheads). In the STI-STAR susceptibility maps, veins often appeared less well defined. Additionally, in some of the susceptibility maps reconstructed with both STI-based methods, the susceptibility in specific veins or venous segments, e.g., within the SSS or StrS, appeared lower than in the surrounding brain tissue (Figure 3, arrows), resulting in an overall reduced conspicuity of these veins.

**Figure 2:**
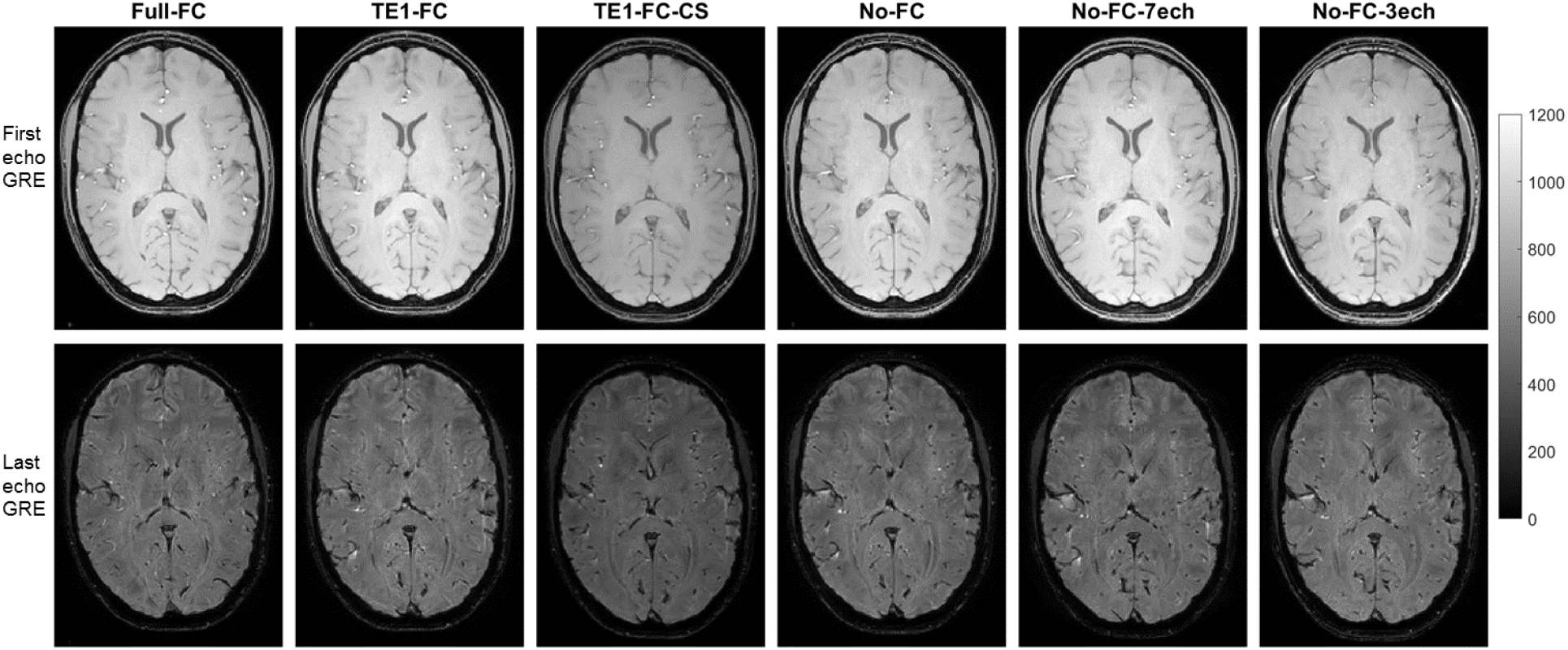
Representative GRE magnitude images of the first (top row) and last (bottom row) echo for each of the six sequences. Similar slices were selected for each sequence. The image quality was comparable across all multi-echo GRE imaging sequences. All images are scaled in arbitrary units. The image brightness can be affected by automatic adjustment of the scanner.

**Figure 3:**
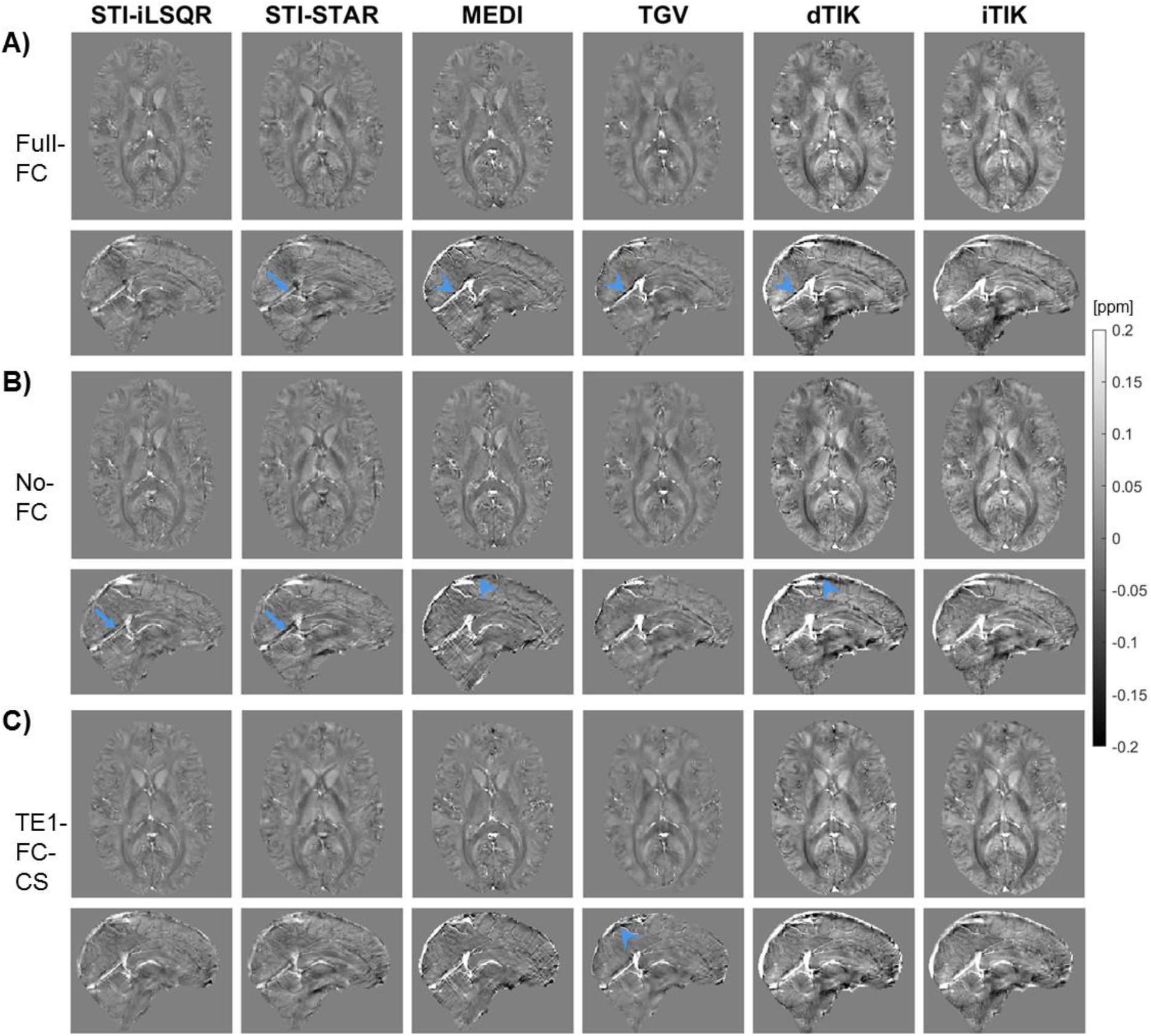
Representative transverse and sagittal slices are shown for all susceptibility maps reconstructed with the six different QSM pipelines (columns) for a healthy subject. Similar transverse and sagittal slices were selected for (A) Full-FC, (B) No-FC, and (C) TE1-FC-CS sequences. Differences in the susceptibility maps can be seen between the six reconstruction methods including the degree of SSS erosion, substantially decreased susceptibility values around veins (arrowheads), and reduced susceptibility values within venous areas (arrows).

### 3.2. Performance of automated whole-brain vessel segmentation

For the same acquisition sequence, the number and location of automatically segmented venous voxels differed across QSM reconstructions (Figure 4A). While about 10%-25% of automatically MVF-based segmented venous voxels were recognized by the MVF algorithm in all six susceptibility maps from the same sequence (Figure 4A, blue and yellow arrows), others were only segmented in some or just a single QSM reconstruction (Figure 4A, arrowheads). The STI-based and MEDI reconstructions had the highest numbers of uniquely segmented venous voxels, i.e., voxels that were only segmented in one QSM map from a single sequence (about 3000 to 7200 voxels), whereas susceptibility maps calculated using both TIK-based methods had the lowest numbers of uniquely segmented voxels (about 950 voxels, Figure 5A). Similarly, the ratios between the number of uniquely segmented voxels and the number of voxels segmented commonly in all six susceptibility maps from different QSM reconstruction methods were highest for the STI-based and MEDI reconstructions and lowest for TIK-based methods (Figure 5B).

**Figure 4:**
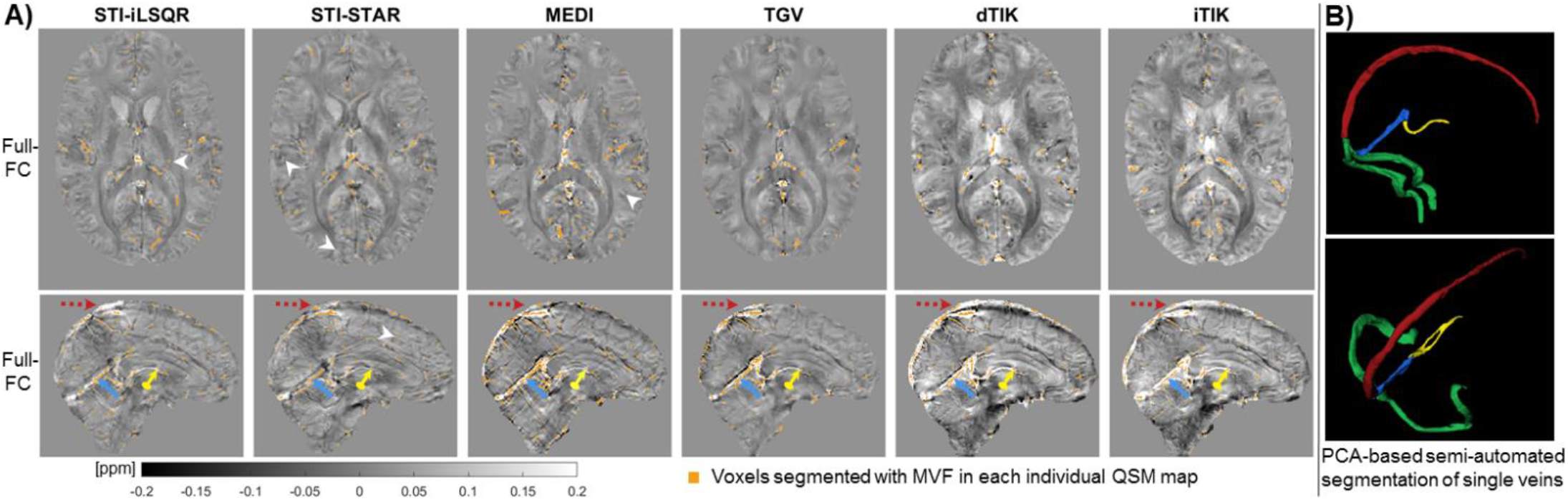
Representative results of vein segmentations for different QSM reconstruction methods for (A) MVF-based automated whole-brain segmentations and (B) PCA-based semi-automated segmentation of single representative veins. In (A), the same transverse and sagittal slices are shown for each QSM reconstruction (columns) of the Full-FC data with the segmented voxels overlaid (orange). The arrows point to the superior sagittal sinus (red dotted arrows), the straight sinus (blue arrows), and an internal cerebral vein (yellow circle-ended arrow). White arrowheads point to voxels uniquely segmented in the corresponding susceptibility map. In (B), the superior sagittal sinus (SSS), the straight sinus (StrS), both transverse sinuses (TraS), and the internal cerebral veins (ICVs) are colored in red, blue, green, and yellow, respectively.

**Figure 5:**
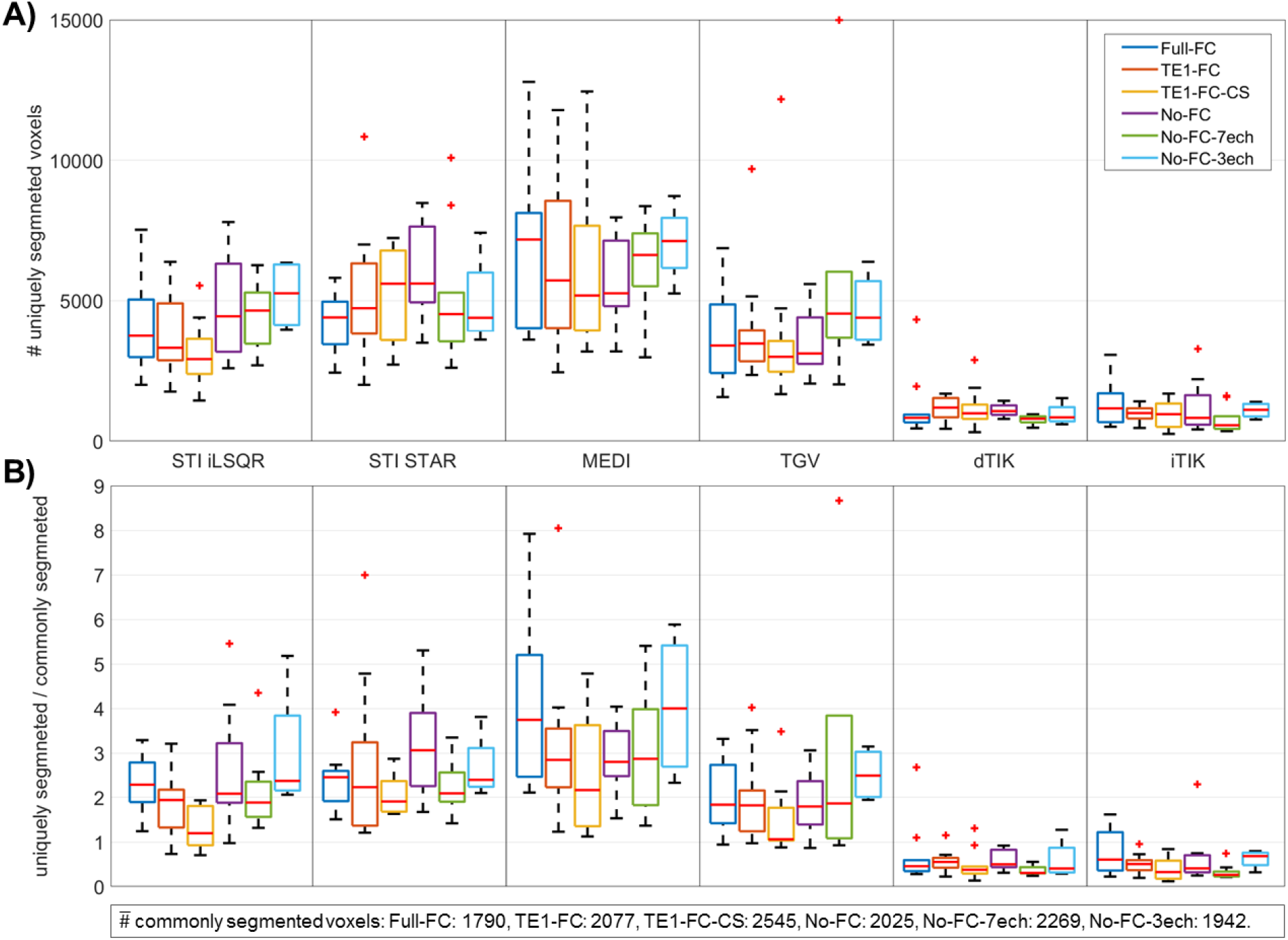
Relative agreement between automated whole-brain segmentations across sequences and processing pipelines. For each QSM method and acquisition sequence, panel (A) shows the number of voxels uniquely segmented by the MVF algorithm for that combination of method and sequence. In panel (B), the number of uniquely segmented voxels derived from the same acquisition sequence is shown normalized relative to the number of voxels commonly segmented by all QSM reconstruction methods for that sequence. For each sequence, the number of commonly segmented voxels averaged across subjects is shown at the bottom of the figure. Each acquisition sequence is represented using a different color and the six QSM methods are grouped by column. In both panels, the boxplots represent distributions across subjects.

Dice scores quantifying the relative overlap between whole-brain segmentations are reported in Supplementary Figure S2. For all imaging sequences, the highest Dice scores (range: 0.5-0.7) were found between the two TIK-based susceptibility maps, whereas the lowest Dice scores (range: 0.2-0.45) were found between STI-STAR and all non-STI-based pipelines (Supplementary Figure S2).

Automated whole-brain segmentations were compared to independent PCA-based semi-automated segmentations of single representative veins (Figure 4B). MEDI and TGV reconstructions had the highest fractions of voxels correctly segmented within the representative veins (Figure 6). Supplementary Figure S3 shows the fraction of connected voxels in the MVF-based segmentations within the corresponding semi-automated ICVs segmentation. STI-STAR reconstructions had the lowest fractions of connected voxels (average: 7%-9%), STI-iLSQR and both Tikhonov-based reconstructions had a slightly higher connected fraction (average: 9%-28%), whereas MEDI and TGV reconstructions had the highest connected fraction (average: 22%-35%).

**Figure 6:**
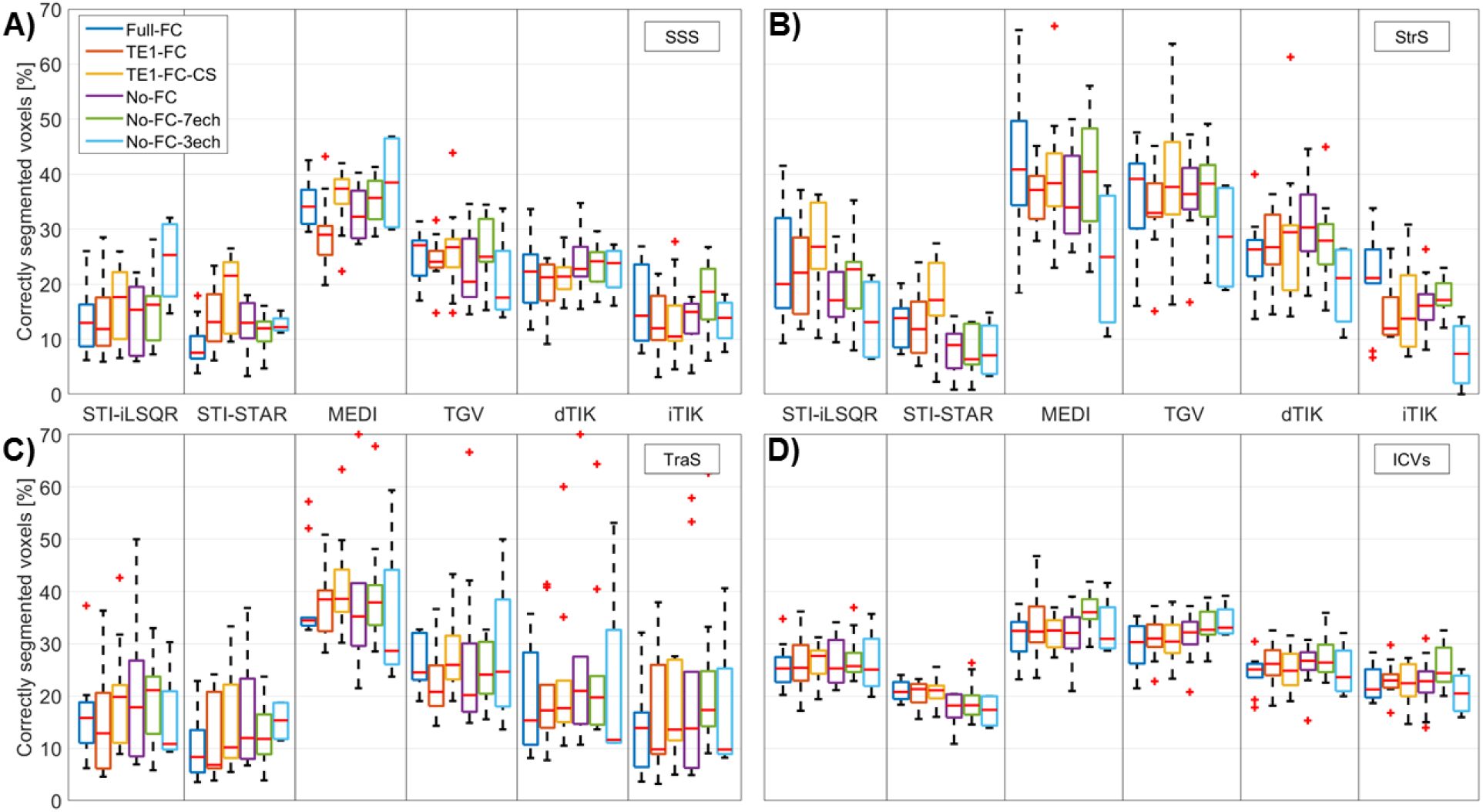
Boxplots of the fraction of PCA-segmented voxels segmented correctly by the MVF algorithm within (A) the superior sagittal sinus (SSS), (B) the straight sinus (StrS), (C) both transverse sinuses (TraS), and (D) the internal cerebral veins (ICVs). The four different (bilateral) vein segmentations were obtained using semi-automated segmentation on PCA images and used as a gold standard after co-registration to each individual susceptibility map. Each acquisition sequence is represented using a different color and the six QSM methods are grouped by column. In all four panels, the boxplots represent distributions across subjects.

Similar results were obtained from automated whole-brain vessel segmentation of the pilot study data (including five different subjects imaged on another scanner model from the same vendor). The numbers of uniquely segmented venous voxels and the ratios between uniquely segmented and commonly segmented voxels (Supplementary Figure S4) were again highest for MEDI and lowest for TIK-based reconstructions. Furthermore, Dice scores between pairs of QSM methods in the pilot study data (Supplementary Figure S5) correlated well with the corresponding Dice scores of the main study (Supplementary Figure S2). Analyses requiring PCA for segmentation could not be performed on the pilot study data, since PCA images were only acquired in three of the five subjects due to time constraints, resulting in a sample size too small for meaningful statistical analysis.

### 3.3. Susceptibility quantification inside venous segmentations

In all subjects, STI-based reconstructions yielded the lowest average susceptibility values for both the whole-brain automated vein segmentations (Figure 7A) and the single-vein semi-automated segmentations (Figure 8). Notably, TIK-based versus STI-based reconstructions yielded similar whole-brain venous densities (∼0.7%-1.0%), but consistently higher average susceptibility values (Figure 7). MEDI and TGV reconstructions resulted in slightly higher venous densities in the brain (∼1.0%-1.4%) (Figure 7B). In all venous segmentations, dTIK reconstructions resulted in the highest venous susceptibility values and thereby in the lowest SvO_2_ values (Supplementary Table S6), but also had the most variable venous susceptibility values (Figures 7A and 8).

**Figure 7:**
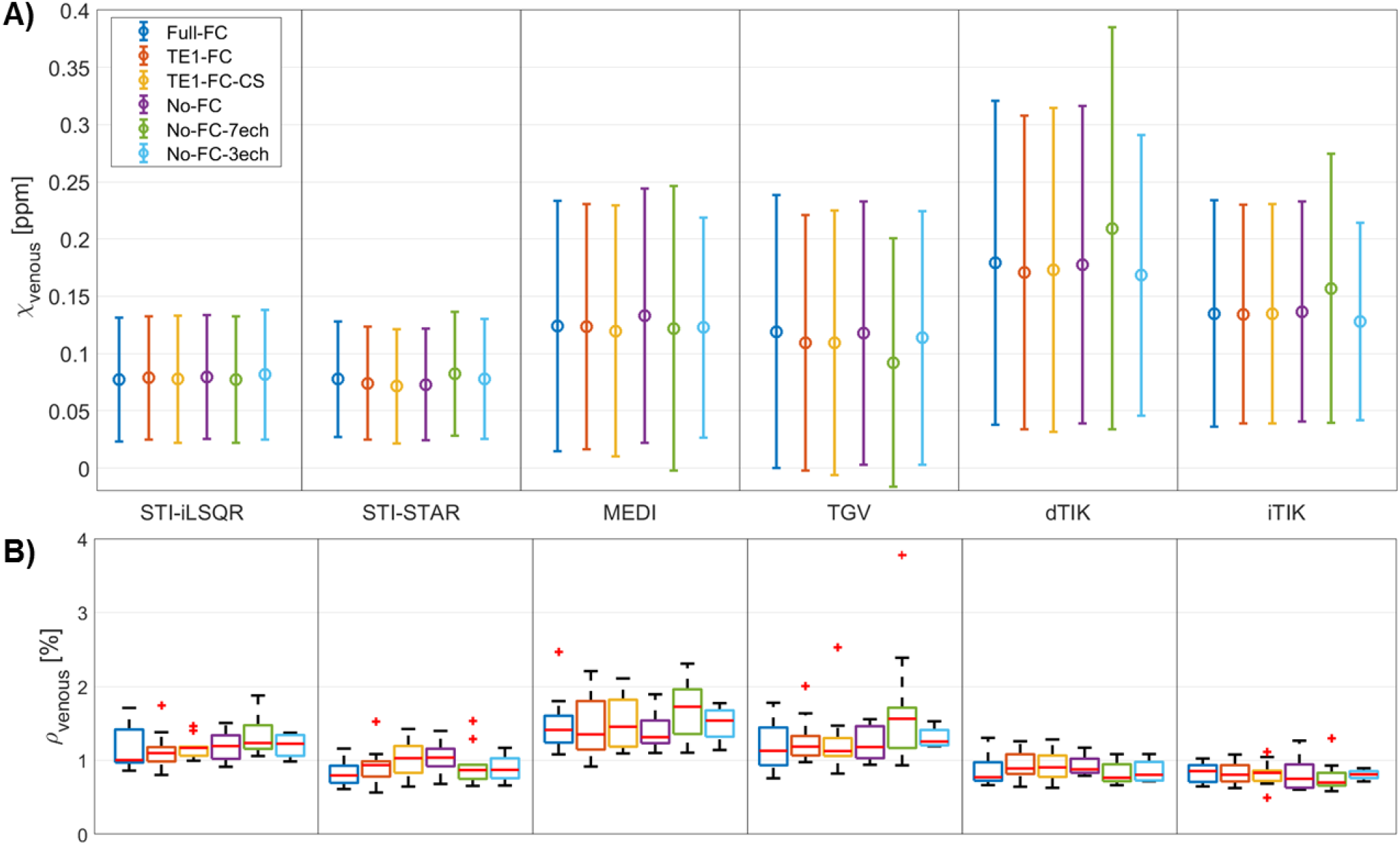
Whole-brain (A) mean and standard deviations of venous susceptibility values across subjects and (B) subject-mean venous densities for the six acquisition sequences (different colors) and six QSM methods (columns). Venous susceptibility and venous density values were calculated across all voxels obtained from multiscale vessel filtering on individual susceptibility maps within a common minimum-size brain mask. Differences in mean venous susceptibility and subject-mean venous density are greater for different QSM reconstruction methods than for different acquisition settings.

**Figure 8:**
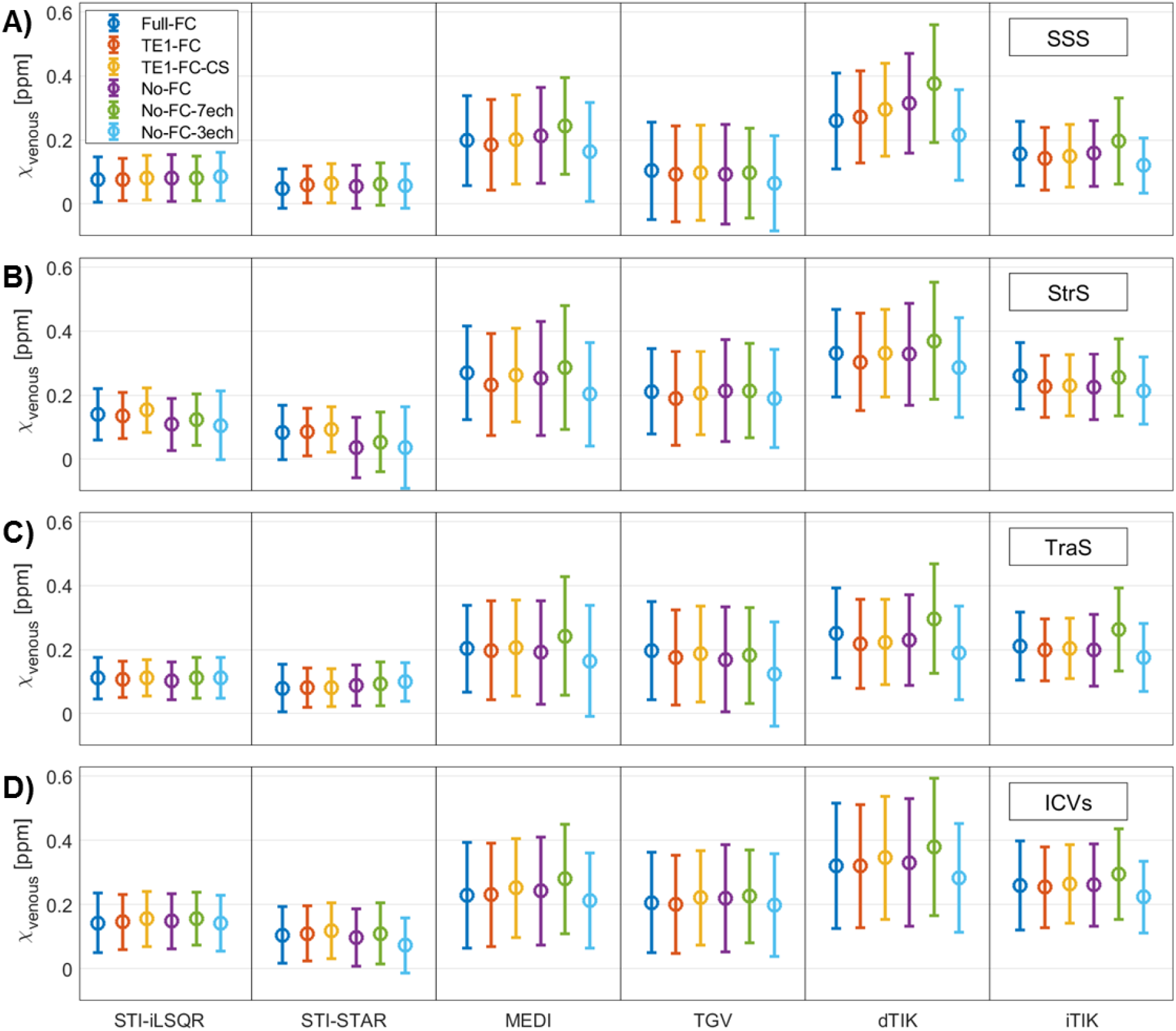
Mean and standard deviations of venous susceptibility values in single-vein segmentations of (A) the superior sagittal sinus (SSS), (B) the straight sinus (StrS), (C) both transverse sinuses (TraS), and (D) the internal cerebral veins (ICVs). Data are shown for the six acquisition sequences (different colors) and six QSM methods (columns). Mean venous susceptibility was calculated across all voxels obtained from semi-automated segmentation on PCA images and co-registered to each individual susceptibility map. Differences in mean venous susceptibility are greater for different QSM reconstruction methods than for different acquisition settings.

Similar average susceptibility values and venous densities (∼0.6%-1.3%) were obtained from five different subjects acquired on another scanner model from the same vendor (Supplementary Figure S7).

Two-way repeated measures ANOVA with Greenhouse-Geisser correction for non-sphericity revealed that significant differences in the average whole-brain venous susceptibility were linked to both the type of imaging sequence (F(4, 36) = 4.19, p =0.007, η^2^ = 0.006) and the QSM reconstruction method employed (F(2.36, 20.32) = 222.10, p < 0.001, η^2^ = 0.866). Additionally, there was a statistically significant interaction between the effects of imaging sequence and QSM method on venous susceptibility (F(5.38, 48.38) = 5.13, p = 0.001, η^2^ = 0.028). Similar results were found for the average susceptibility in single-vein segmentations (Supplementary Table S8).

Paired samples t-tests with Benjamini-Hochberg correction for a false discovery rate revealed that most average venous susceptibility values calculated with different QSM methods were significantly different (Supplementary Table S9). However, for various acquisition sequences and across several venous segments, no significant differences in average venous susceptibility were found between iTIK and MEDI as well as between iTIK and TGV. Furthermore, some pairs of reconstruction methods did not show statistically significantly different mean susceptibility values only within single venous segments, such as STI-iLSQR and TGV within the SSS or MEDI and dTIK within the TraS segmentations.

Much smaller differences in venous susceptibility values were found between data acquired with different imaging sequences than across the six QSM methods (Figures 7A and 8). However, venous susceptibility values acquired with the No-FC-7ech sequence were frequently slightly increased compared to other sequences. Similarly, paired samples t-tests with Benjamini-Hochberg correction comparing mean susceptibilities from different acquisition sequences revealed only a few statistically significant differences (Supplementary Table S10). These differences were mainly found between the No-FC-7ech sequence and any of the other sequences.

For all single-vein segmentations, 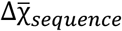, the maximal difference between the six sequence-mean venous susceptibility values 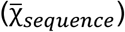 was almost constant (varying between 0.05 and 0.06 ppm), while 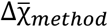 (the maximal difference between the six method-mean venous susceptibility values 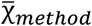) varied more strongly across the segmentations (between 0.14 and 0.26 ppm) (Table 2). These results show that the influence of the QSM reconstruction method on the average venous susceptibility values was much stronger than the influence of the acquisition sequence, ranging from a factor of 2.7 (TraS) to 5.8 (StrS). Whole-brain segmentations, which contained many more voxels compared to single-vein segmentations, resulted in smaller maximal differences between sequence-mean and method-mean susceptibilities. However, in the whole-brain ROIs, the influence of the QSM reconstruction method on the average venous susceptibility was 11.6 times larger than the influence of the sequence (Table 2), which concurs with a factor of 12.4 obtained for the pilot study data acquired on a different scanner model (Supplementary Table S11).

**Table 2:** Maximal differences between the method-mean 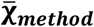 and sequence-mean 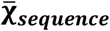 venous susceptibility values and their quotient. The method-mean values were calculated by averaging over venous susceptibility values acquired with six different sequences but reconstructed with the same QSM method. Accordingly, sequence-mean values were calculated by averaging over venous susceptibility values acquired with the same sequence but reconstructed with six different QSM methods.

| Maximal average differences in $\bar{\chi}$-venous | MVF | SSS | StrS | TraS | ICVs |
| --- | --- | --- | --- | --- | --- |
| $\Delta\bar{\chi}_{sequence}$ [ppm] | 0.009 | 0.058 | 0.045 | 0.054 | 0.053 |
| $\Delta\bar{\chi}_{method}$ [ppm] | 0.104 | 0.236 | 0.263 | 0.143 | 0.231 |
| $\Delta\bar{\chi}_{method} / \Delta\bar{\chi}_{sequence}$ | 11.56 | 4.07 | 5.84 | 2.65 | 4.36 |

## 4. Discussion

This study aimed to compare acquisition sequences and QSM reconstruction pipelines for venous QSM. The effect of flow compensation during image acquisition was evaluated by applying sequences incorporating different flow compensation schemes versus no flow compensation when using monopolar readout gradients. The effect of using different QSM reconstruction methods was evaluated by testing pipelines from popular QSM toolboxes compared to pipelines previously optimized for venous QSM. The effects of acquisition sequence and QSM method on the accuracy of venous susceptibility values were assessed by applying all combinations of sequences and QSM pipelines to the multi-echo GRE data acquired in ten healthy subjects.

### 4.1. Influence of the imaging sequence on venous QSM

Generally, there was a high similarity in the visual appearance of susceptibility maps, venous susceptibility values, and the performance of whole-brain segmentation between acquisition sequences. Slight differences were only found for the two sequences without flow compensation (No-FC-3ech and No-FC-7ech) that had a different number of echoes from all other sequences (which had four echoes). Susceptibility maps reconstructed from the No-FC-3ech sequence resulted in slightly lower average venous susceptibility values within single-vein segmentations (Figure 8) and in slightly lower variances in susceptibility in the whole-brain segmentations (for the MEDI, dTIK, and iTIK pipelines) (Figure 7A). The smaller sample size acquired with this sequence (four subjects) did not allow a direct comparison with other sequences (ten subjects). However, it appears that, for accurate multi-echo venous QSM, maximizing the SNR by using the minimum number of TEs required for multi-echo fitting using the MEDI toolbox as in No-FC-3ech had a slightly worse performance than sequences with more echoes.

Sequences with four echoes had more consistent mean susceptibility values and it is possible that including more echoes in the nonlinear fit (used for all but the STI-based methods) slightly improved the accuracy of venous QSM. The No-FC-7ech sequence resulted in slightly higher venous susceptibility values compared to all other sequences (Figures 7A and 8). Here, a possible cause for measuring higher venous susceptibility values may be phase accumulation due to the lack of flow compensation. Moreover, compared to the two other sequences without any type of flow compensation, the No-FC-7ech sequence had a lower signal-to-noise ratio (due to its higher bandwidth). Venous susceptibility values differed least between the fully flow compensated sequence (Full-FC) and the sequences with only first-echo flow compensation (TE1-FC and TE1-FC-CS) or the sequence without flow compensation and matched echo times (No-FC). This suggests that the number of echoes had more of an effect on the susceptibility values than the flow compensation scheme.

Generally, different flow compensation implementations yielded comparable venous susceptibility values. It is possible that the monopolar readout gradients employed in all imaging sequences mitigated the lack of flow compensation by effectively applying a partial form of flow compensation, even when full flow compensation was not explicitly implemented. Therefore, sequences without flow compensation or with only first-echo flow compensation appear as suitable for quantitative analyses as sequences with full flow compensation when monopolar read gradients are used.

No significant differences in image quality and venous susceptibility were detected when comparing the Compressed SENSE-accelerated sequence (TE1-FC-CS) with the other SENSE-accelerated sequences. Furthermore, the TE1-FC-CS sequence enabled reliable whole-brain segmentations, resulting in the lowest ratios of uniquely segmented voxels to voxels commonly segmented in all six susceptibility maps from different QSM reconstruction methods compared to all other sequences (Figure 5B). Therefore, our results suggest that, for venous QSM, Compressed SENSE could be used to obtain clinically feasible scan times (5-6 minutes) without introducing significant venous susceptibility differences compared to SENSE accelerated sequences.

### 4.2. Influence of the QSM reconstruction pipeline on venous QSM

Our results show that the choice of QSM processing pipeline is crucial for correctly reconstructing high susceptibility values in veins. The QSM method had a much (2.7 to 11.6 times) greater effect on the average venous susceptibility than the acquisition sequence in general or the use of flow compensation in particular (Table 2, Figures 7A and 8). These findings were reproduced using data that were acquired on a different scanner model from the same vendor (Supplementary Table S11 and Supplementary Figure S7). Furthermore, different vein segmentation approaches did not influence our results: similar trends were seen when using either automated whole-brain or semi-automated single-vein segmentation methods (Figures 7A and 8).

### 4.3. Accuracy of automatic whole-brain segmentations

High fractions of correctly segmented voxels within the gold-standard single-vein segmentations (Figure 6) and high fractions of connected voxels within the ICVs (Supplementary Figure S3) were considered to indicate accurate segmentation performance. Additionally, the number of uniquely segmented voxels (Figure 5A) and the ratio between the number of uniquely segmented voxels and the number of voxels commonly segmented in all six susceptibility maps from different QSM reconstruction methods (Figure 5B) can also be used to provide information on the agreement between and performance of automated whole-brain segmentations. Automated whole-brain segmentations based on susceptibility maps from both STI-based reconstructions resulted in the lowest fractions of correctly segmented voxels. Additionally, STI-STAR yielded the lowest fraction of connected voxels within the ICVs and high numbers of uniquely segmented voxels. These results combined indicate both a large number of false positive and false negative voxels in the automatic segmentations for STI-STAR, suggesting that STI-STAR may not yield accurate automatic vein segmentations using the MVF algorithm. TIK-based methods yielded higher quality automated whole-brain segmentations, resulting in a low number of uniquely segmented voxels, a low ratio between uniquely and commonly segmented voxels, and slightly increased fractions of connected voxels compared to STI-STAR. Additionally, TIK-based methods showed the smallest standard deviations of ratios between uniquely and commonly segmented voxels indicating the most stable performance across subjects. The highest ratio between uniquely and commonly segmented voxels and the largest variability across subjects and across sequences was found in susceptibility maps reconstructed with MEDI (Figure 5). Furthermore, the highest fractions of connected voxels within an independently segmented representative vein and of correctly segmented voxels within several representative veins were found for MEDI and TGV. Together, these results suggest that MEDI may provide better performance in the segmentations of the representative veins, resulting in a higher number of correctly segmented voxels compared to other methods. Overall, susceptibility maps from MEDI and TGV reconstructions did not appear optimal for general brain QSM because of streaking artifacts and rather low gray/white matter contrast, but for automated vein segmentation with the MVF algorithm, they appeared to provide the most accurate results among the investigated methods.

### 4.4. Differences between processing pipelines

Susceptibility maps reconstructed using methods from the STI Suite were found to differ most from the other QSM reconstructions, e.g., resulting in comparatively low venous susceptibility values (Figure 7A and 8) and low fractions of correctly segmented voxels (Figure 6). These differences between susceptibility maps calculated using STI-based pipelines and those reconstructed using all other pipelines could be caused by differences in multi-echo GRE signal combination. For the STI-based pipelines, the phase at each echo was first unwrapped and then averaged over all echoes using a TE-based weighting approach (Li et al., 2015; Wei et al., 2015). Conversely, all other pipelines combined the complex GRE signal by nonlinear fitting over TEs before performing spatial phase unwrapping (Liu et al., 2013). Unlike fitting over TEs, averaging over TEs does not allow removal of phase offsets from the fitted field map. Moreover, Laplacian-based phase unwrapping could alter the linearity of the signal phase over time by inherently performing some degree of background field suppression, thus yielding inaccurate values if the phase is combined over TEs after Laplacian unwrapping (Biondetti et al., 2016; Schweser et al., 2013). These two aspects could have jointly contributed to the generally lower accuracy of STI-based venous QSM. Additionally, in the STI-STAR pipeline, high-susceptibility areas are reconstructed separately from the surrounding tissue by applying a two-step algorithm (Wei et al., 2015). In the first step, the regularization parameter is tuned to “high-susceptibility sources”, which are reconstructed separately, and then the forward model (Marques and Bowtell, 2005) is applied to estimate and remove field variations from these sources from the original field map. In the second step, “low-susceptibility sources” are reconstructed in the rest of the brain by tuning the regularization parameter to brain parenchyma (Wei et al., 2015). However, the first regularization parameter was originally tuned for detecting hemorrhages, which are characterized by much higher susceptibility values (up to ∼1 ppm) than normal appearing veins (∼0.3-0.4 ppm) (Wei et al., 2015). Thus, in our study, it is possible that voxels within a normal venous susceptibility range were largely omitted in the first step of the STI-STAR method, but then over-regularized in the second step, resulting in unrealistically low venous susceptibilities. Note that tuning the regularization parameters to different values was not possible because the corresponding function in the STI toolbox is designated as private.

Generally, differences between QSM methods in the regularization used in the local field-to-susceptibility inversion step can influence the appearance of susceptibility maps and the variability of susceptibility values. It is possible that a lower degree of regularization could explain the higher variance of venous susceptibility found for dTIK and the lower overall homogeneity generally seen in TIK-based susceptibility maps possibly caused by residual background fields.

### 4.5. Quantification of venous oxygenation

In most of the combinations of acquisition sequence and QSM reconstruction method, the venous susceptibility tended to be underestimated and consequently the resulting SvO_2_ was overestimated compared to literature values (Biondetti et al., 2019; Fan et al., 2014; Xu et al., 2014). Whole-brain SvO_2_ was overestimated even using MEDI and TGV, which seemed promising for automatically quantifying whole-brain venous oxygenation since they provided the most accurate automatic segmentation results. QSM reconstructions from dTIK yielded venous oxygenation values (Supplementary Table S6) closest to venous oxygenation obtained by gold-standard PET measurements of the oxygen extraction fraction (OEF, SvO_2_ = 1 – OEF). In all single-vein segmentations, the average venous susceptibility in dTIK QSM ranged between 0.19 ppm and 0.38 ppm corresponding to an SvO_2_ of 74.9%-60.9%, which is in good agreement with literature values measured using ^15^O PET (i.e., 59%-72%) (Cho et al., 2020; Ibaraki et al., 2008; Ishii et al., 1996) and MRI based on (apparent) transverse relaxation times (T2* and T2) (i.e., 63%-70.6%) (Barhoum et al., 2015; Cho et al., 2020; Fan et al., 2016; Jain et al., 2013). In MEDI and iTIK reconstructions, accurate SvO_2_ values were only found in the StrS and ICVs segmentations, whereas SvO_2_ values in the SSS and TraS were overestimated.

Generally, whole-brain venous susceptibility distributions within dTIK reconstructions had the highest variance (Figure 7A), suggesting less efficient noise or streaking artifact reduction compared to other pipelines. Nevertheless, the good correspondence of SvO_2_ from dTIK (and partly also from iTIK or MEDI) QSM with SvO_2_ measured using ^15^O PET suggests that QSM maps with the best overall visual appearance might not provide the most accurate susceptibility quantification (and SvO2 measurement) in veins. Similarly, the QSM reconstruction challenge 2.0 showed that errors in the estimation of susceptibility values in the veins were particularly high, even for many of the top-ranked algorithms (Bilgic et al., 2020). These findings indicate that the quantification of susceptibilities in venous blood remains prone to errors and needs further optimization.

### 4.6. Limitations and recommendations for future studies

To investigate differences in susceptibility values within the same venous voxels, all evaluations were performed using a minimum-size brain mask, calculated as the intersection of all QSM-based brain masks. However, after applying this combined brain mask, the semi-automated SSS and TraS segmentations were significantly reduced in size and a large portion of these superficial vessels was excluded from both the whole-brain and the single-vein analyses. This was caused mainly by MEDI and TGV reconstructions eroding larger numbers of superficial voxels compared to other methods. When designing the image reconstruction pipeline, future studies for venous QSM should account for the degree of brain erosion. For venous QSM, further work is needed to evaluate the accuracy of methods that do not require brain erosion during background field removal (Liu et al., 2017) and could thus enable segmentation of the entire venous vasculature.

Partial volume effects of vessels with the surrounding brain tissue is a common confounding factor in venous QSM. This problem was addressed automatically for MVF-based whole-brain segmentations through a model-based approach (Bazin et al., 2015) and using erosion for single-vein segmentations. However, in smaller veins, erosion was not feasible since it would have removed most segmented venous voxels. Thus, our single-vein analyses were mainly limited to large dural sinuses and one ROI drawn on smaller veins (the ICVs) that was analyzed without applying erosion.

In this study, image acquisition was limited to a single vendor and one field strength. Future studies on 3 T systems from different vendors are needed to evaluate, for example, the feasibility of multi-echo GRE MRI using compressed sensing. Moreover, since T_2_* values shorten with increasing field strength and the optimal phase SNR for a given tissue is achieved when TE equals T_2_* of that tissue (Wu et al., 2012a), future studies at ultra-high fields (e.g., 7 T) could investigate the potential reduction in acquisition time achievable by reaching the desired phase SNR at shorter TEs.

## 5. Conclusion

To conclude, the effect of different QSM reconstruction methods on mean venous susceptibility values was several times greater than the effect of varying acquisition settings including flow compensation. This indicates that specific optimization of QSM algorithms and pipelines is essential for accurate venous QSM and, in turn, to enable reproducible venous susceptibility quantification across studies. Future venous QSM studies need to carefully select their QSM reconstruction method based on their individual research question and to consider that the high susceptibilities in veins could most accurately be quantified by QSM reconstruction methods that do not necessarily provide the “best-looking” maps in other tissue types.

## Acknowledgements

We thank Dr. Guillaume Gilbert (MR Clinical Science, Philips Healthcare, Markham, Canada) for providing the software for full multi-echo flow compensation and Dr. Anita Karsa (Department of Medical Physics and Biomedical Engineering, University College London (UCL), London, UK) for discussion on implementing the iterative Tikhonov method. We thank Dr. Jakob Meineke (Philips Research, Hamburg, Germany) and Dr. Andreas Hock (Philips Healthcare, Hamburg, Germany) for their support and discussion during preparatory test scans and analyses at the Technical University of Munich. We also thank Dr. Marco Battiston, Dr. Francesco Grussu, and Dr. Marios Yiannakas (NMR Research Unit, Queen Square Multiple Sclerosis Centre, UCL Queen Square Institute of Neurology, UCL, London, UK) and Dr. Rosa Cortese and Dr. Floriana De Angelis (Queen Square MS Centre, UCL Queen Square Institute of Neurology, UCL, London, UK) for their help during the MRI scans at UCL. We are indebted to Prof. Claudia Gandini Wheeler-Kingshott (NMR Research Unit, Queen Square MS Centre, UCL Queen Square Institute of Neurology, UCL, London, UK) for her support with the MRI experiments performed at the Queen Square MS Centre, UCL.

## List of abbreviations

χ: magnetic susceptibility;
ANOVA: analysis of variance;
CS: Compressed SENSE;
dTIK: direct Tikhonov;
GRE: gradient-recalled echo;
ICVs: internal cerebral veins;
iLSQR: iterative least-squares;
iTIK: iterative Tikhonov;
MEDI: Morphology Enabled Dipole Inversion;
MVF: multiscale vessel filtering;
OEF: oxygen extraction fraction;
PCA: phase contrast angiography;
PE: phase encoding;
QSM: quantitative susceptibility mapping;
ROI: region of interest;
SNR: signal-to-noise ratio;
SSS: superior sagittal sinus;
STAR: streaking artifact reduction;
STI: Susceptibility Tensor Imaging;
StrS: straight sinus;
SvO_2_: venous oxygen saturation;
TE: echo time;
TGV: total generalized variation;
TIK: Tikhonov;
TraS: transverse sinuses.

