## Supplemental figures and tables for "Investigating the Effect of Flow Compensation and Quantitative Susceptibility Mapping Method on the Accuracy of Venous Susceptibility Measurement"

### Supplementary material - Venous Quantitative Susceptibility Mapping

| | Full-FC | TE1-FC | TE1-FC-CS | No-FC | No-FC-7ech | No-FC-3ech |
|---|---|---|---|---|---|---|
| $\alpha_{dT}$ | 0.0703 | 0.0715 | 0.0741 | 0.0712 | 0.0739 | 0.0696 |

**Supporting Table S1: Table of sequence-mean regularization parameters $\alpha$ for direct implementations of the Tikhonov-based calculation of susceptibility maps.** The $\alpha$ parameters were calculated for each data set using the L-curve method. The sequence-mean across all data sets acquired with the same imaging protocol was used as the regularization parameter in the dTIK QSM pipeline.

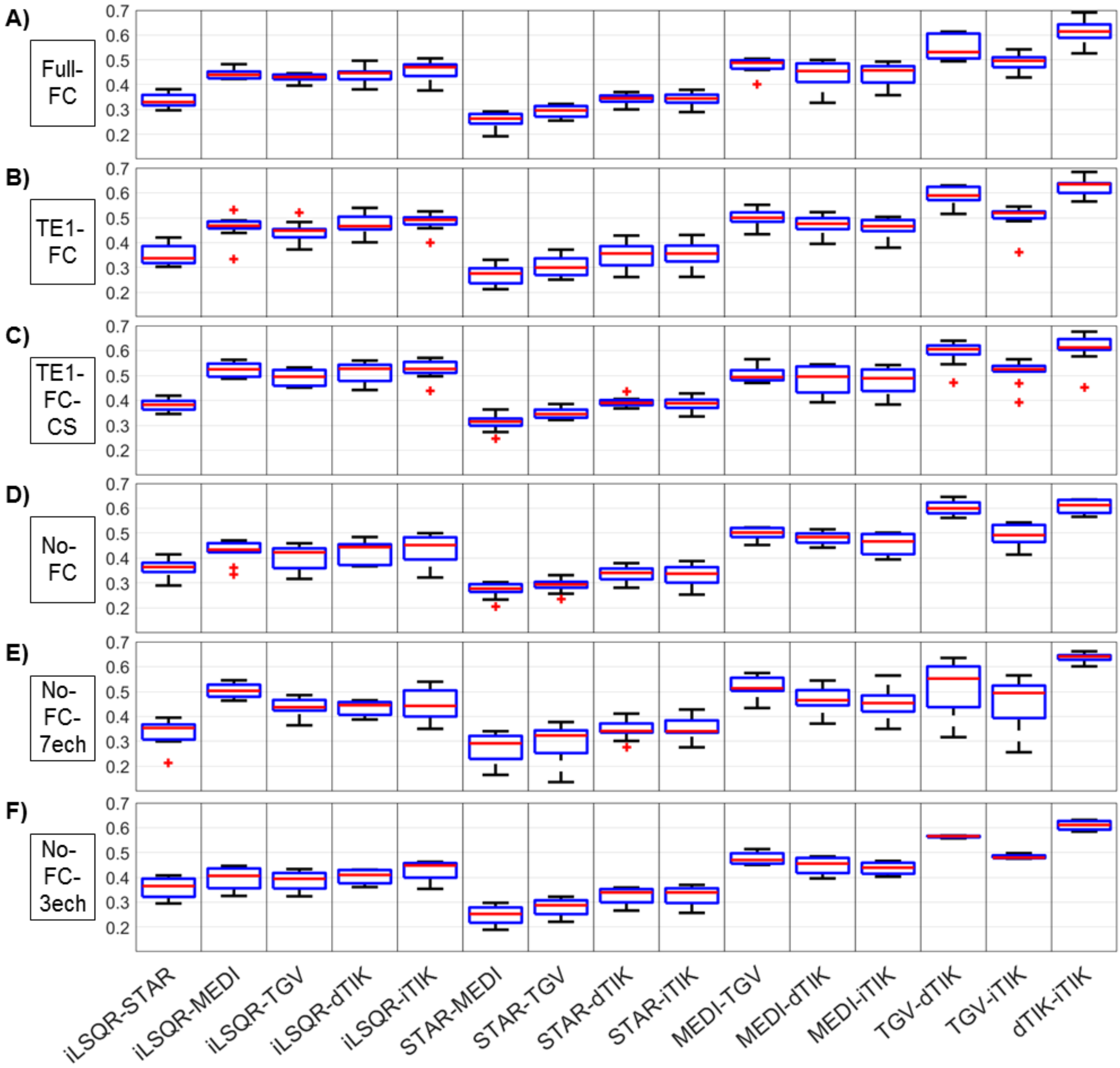


**Supplementary Figure S2: Boxplots of Sørensen-Dice similarity coefficients between whole-brain MVF-based vessel segmentations of susceptibility maps reconstructed with different QSM methods (columns) for each of the six imaging sequences (rows).**

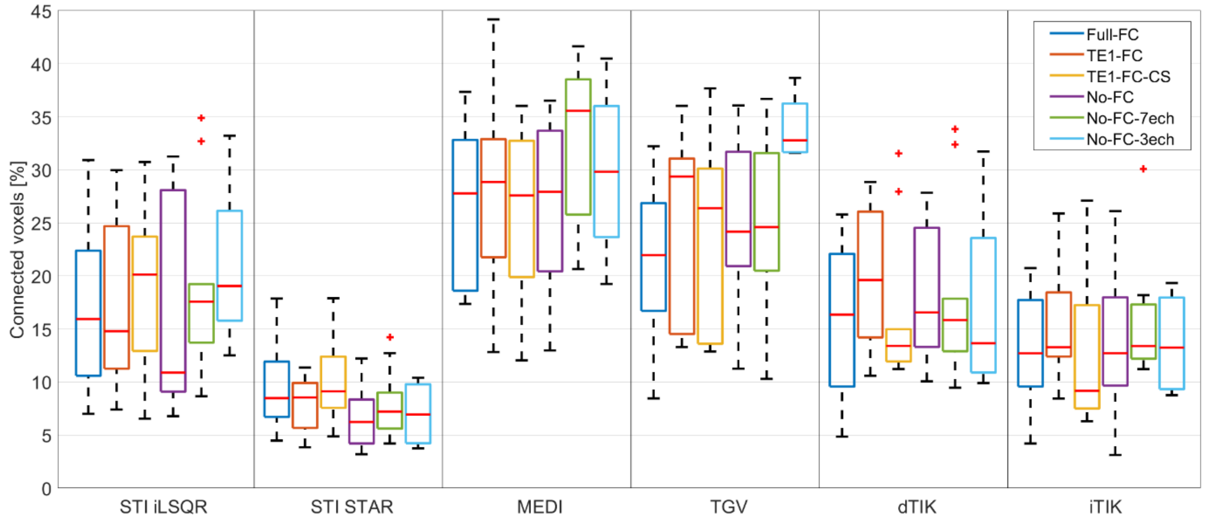


**Supplementary Figure S3: Boxplots of the fraction of voxels segmented by the MVF algorithm that are connected within the PCA-based segmentation of ICVs.** The data are shown for each of the six acquisition sequences (different colors) and six QSM methods (columns).

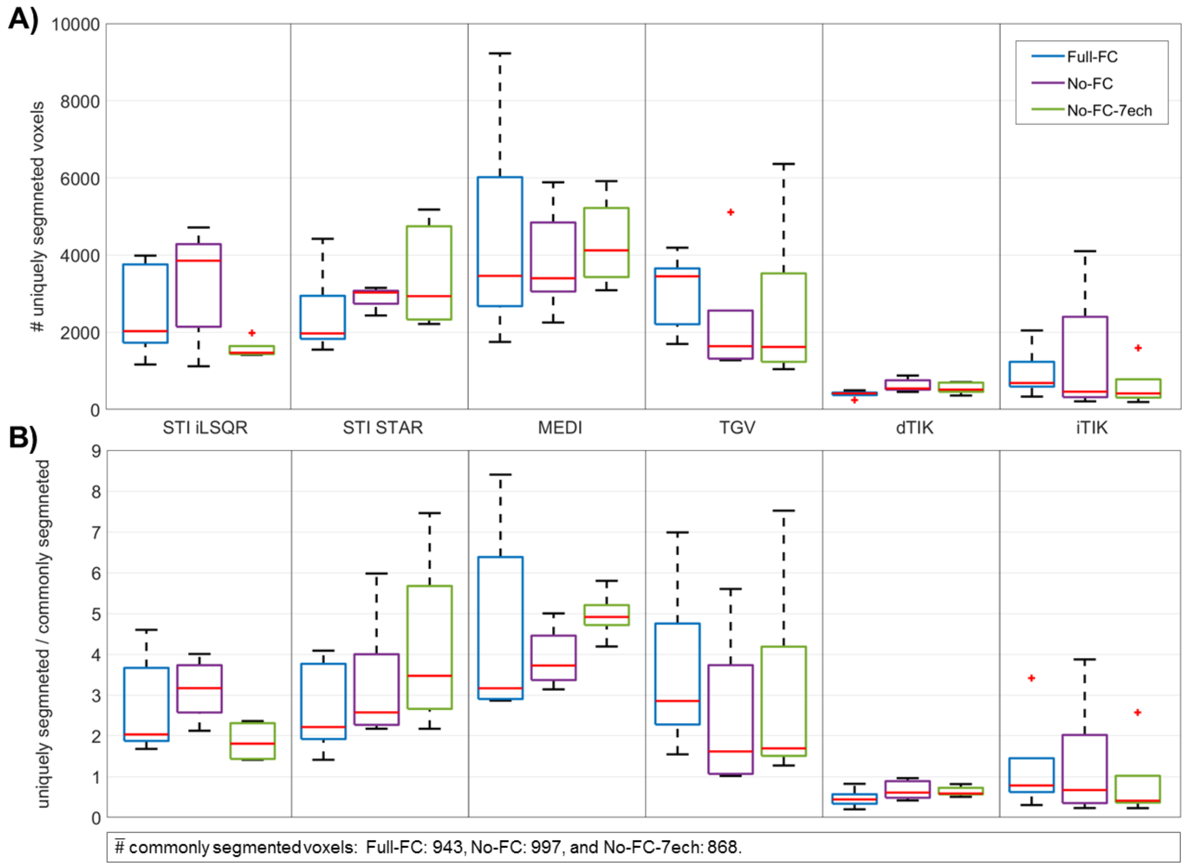


**Supplementary Figure S4: Relative agreement between automated whole-brain segmentations across sequences and processing pipelines from data of the pilot study.** For each QSM method and acquisition sequence, panel (A) shows the number of voxels uniquely segmented by the MVF algorithm for that combination of method and sequence. In panel (B), the number of uniquely segmented voxels derived from the same sequence is shown normalized relative to the number of voxels commonly segmented by all QSM reconstruction methods for that sequence. For each sequence, the number of commonly segmented voxels averaged across the five subjects of the pilot study is shown at the bottom of the figure. Each acquisition sequence is represented using a different color and the six QSM methods are grouped by column. In both panels, the boxplots represent distributions across subjects.

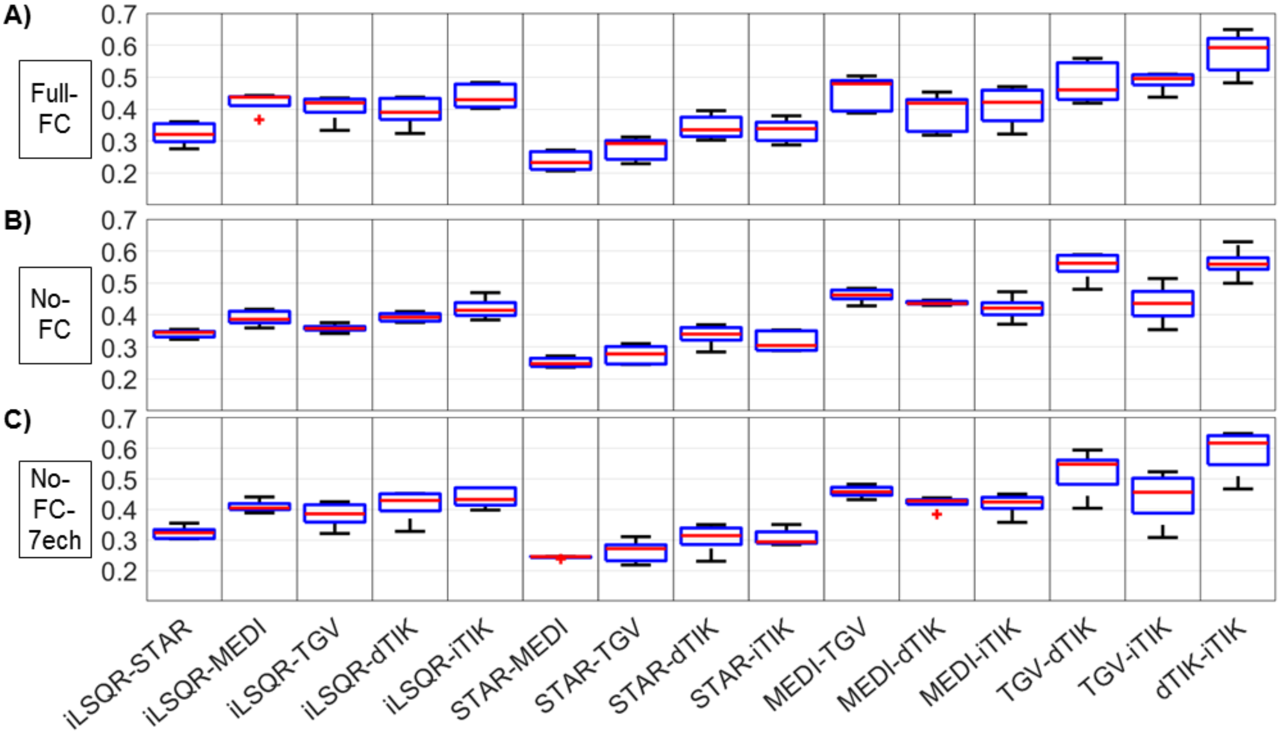


**Supplementary Figure S5: Boxplots of Sørensen-Dice similarity coefficients between whole-brain MVF-based vessel segmentations of susceptibility maps reconstructed with different QSM methods (columns) for each of the three imaging sequences (rows) from data of the pilot study.** Compared to the main study, the data were acquired in five different subjects on another scanner model from the same vendor.

| $SvO_2$ [%] | | iLSQR | STAR | MEDI | TGV | dTIK | iTIK |
|---|---|---|---|---|---|---|---|
| MVF | Full-FC | 83 | 83 | 80 | 80 | 75.6 | 79.3 |
| | TE1-FC | 83 | 83.7 | 80 | 80.8 | 76.4 | 79.3 |
| | TE1-FC-CS | 83 | 83.7 | 80 | 80.8 | 76.4 | 79.3 |
| | No-FC | 83 | 83.7 | 79.3 | 80 | 75.6 | 78.6 |
| | No-FC-7ech | 83 | 83 | 80 | 82.3 | 73.4 | 77.1 |
| | No-FC-3ech | 83 | 83 | 80 | 80.8 | 76.4 | 79.3 |
| SSS | Full-FC | 83 | 85.2 | 74.2 | 81.5 | **69.7** | 77.1 |
| | TE1-FC | 83 | 84.5 | 75.6 | 82.3 | **69** | 78.6 |
| | TE1-FC-CS | 83 | 84.5 | 74.2 | 81.5 | **67.5** | 77.8 |
| | No-FC | 83 | 85.2 | 73.4 | 82.3 | **66** | 77.1 |
| | No-FC-7ech | 83 | 84.5 | **71.2** | 81.5 | **60.9** | 74.2 |
| | No-FC-3ech | 82.3 | 84.5 | 77.1 | 84.5 | 72.7 | 80 |
| StrS | Full-FC | 78.6 | 83 | **69** | 73.4 | **64.6** | **69.7** |
| | TE1-FC | 78.6 | 83 | **71.9** | 74.9 | **66.8** | **71.9** |
| | TE1-FC-CS | 77.8 | 82.3 | **69.7** | 73.4 | **64.6** | **71.9** |
| | No-FC | 80.8 | 85.9 | **70.5** | 73.4 | **64.6** | **71.9** |
| | No-FC-7ech | 80 | 85.2 | **67.5** | 73.4 | **61.6** | **70.5** |
| | No-FC-3ech | 80.8 | 85.9 | 74.2 | 74.9 | **67.5** | 73.4 |
| TraS | Full-FC | 80.8 | 83 | 74.2 | 74.2 | **70.5** | 73.4 |
| | TE1-FC | 80.8 | 83 | 74.2 | 75.6 | 72.7 | 74.2 |
| | TE1-FC-CS | 80.8 | 83 | 74.2 | 74.9 | 72.7 | 74.2 |
| | No-FC | 81.5 | 82.3 | 74.9 | 76.4 | **71.9** | 74.2 |
| | No-FC-7ech | 80.8 | 82.3 | **71.2** | 75.6 | **66.8** | **69.7** |
| | No-FC-3ech | 80.8 | 81.5 | 77.1 | 80 | 74.9 | 75.6 |

| | | | | | | | |
|---|---|---|---|---|---|---|---|
| ICVs | Full-FC | 78.6 | 81.5 | **71.9** | 73.4 | **65.3** | **69.7** |
| | TE1-FC | 77.8 | 80.8 | **71.9** | 74.2 | **65.3** | **70.5** |
| | TE1-FC-CS | 77.1 | 80 | **70.5** | 72.7 | **63.1** | **69.7** |
| | No-FC | 77.8 | 81.5 | **71.2** | 72.7 | **64.6** | **69.7** |
| | No-FC-7ech | 77.1 | 80.8 | **68.3** | **71.9** | **60.9** | **67.5** |
| | No-FC-3ech | 78.6 | 83.7 | 73.4 | 74.2 | **68.3** | 72.7 |

**Supplementary Table S6: Mean values of venous oxygenation ($SvO_2$) within whole-brain and single-vein segmentations from data acquired with different sequences (rows) and reconstructed with various QSM methods (columns).** $SvO_2$ values were calculated from ROI-mean susceptibility values according to Equation 1. $SvO_2$ values that agree with literature values from $^{15}$O PET (i.e., 59%-72%) are marked in bold.

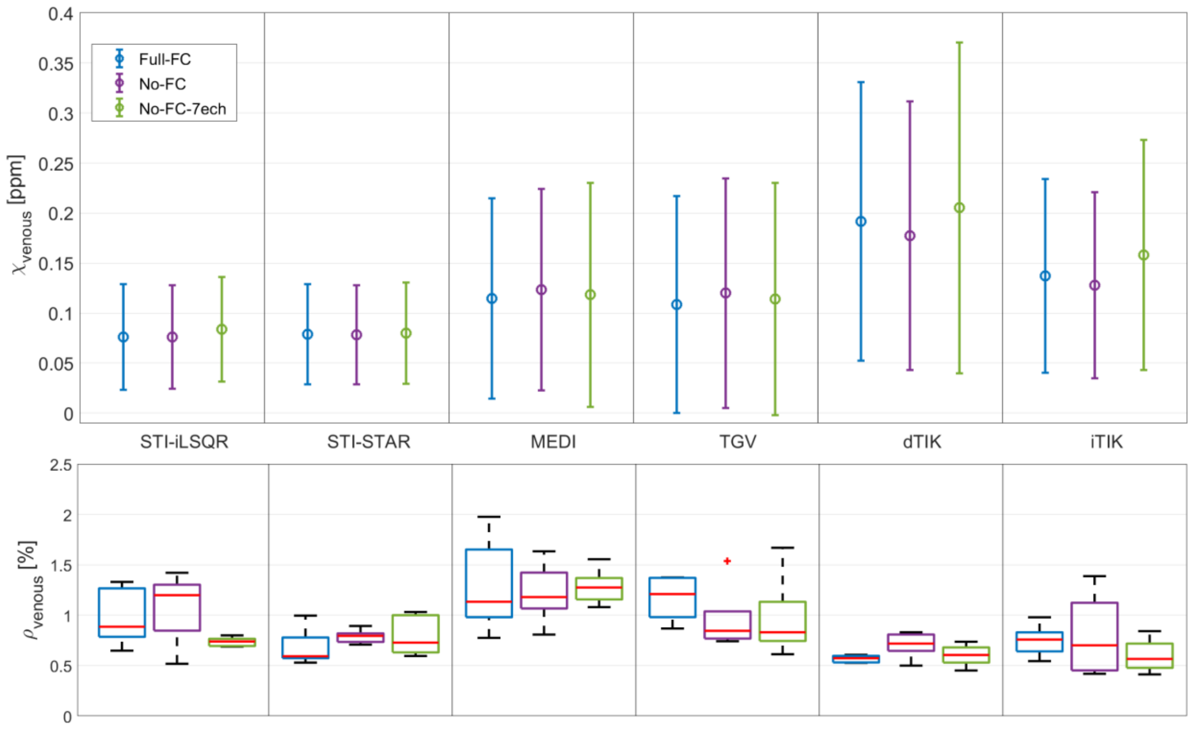


**Supplementary Figure S7: Whole-brain (A) mean and standard deviation of venous susceptibility values across subjects and (B) subject-mean venous densities from data of the pilot study.** In five healthy subjects, three of the sequences of the main study (different colors) were acquired on a different scanner model of the same vendor and processed with six different QSM methods (columns). Venous susceptibility and venous density values were calculated across all voxels obtained from multiscale vessel filtering (MVF) on individual susceptibility maps within a common minimum-size brain mask.

| | Source | Type III Sum of Squares | df | Mean Square | F | Sig. (p-val) | $\eta^2 = \frac{SS_{\text{effect}}}{SS_{\text{total}}}$ |
|---|---|---|---|---|---|---|---|
| **Whole-brain MVF segment.** | Sequence | 0.003 | 4 | 0.001 | 4.189 | 0.007 | **0.0064** |
| | Error (sequence) | 0.007 | 36 | * | | | |
| | Method | 0.408 | 2.258 | 0.181 | 222.100 | * | **0.8662** |
| | Error (method) | 0.017 | 20.323 | 0.001 | | | |
| | Sequence * method | 0.013 | 5.376 | 0.002 | 5.131 | 0.001 | **0.0276** |
| | Error (seq * method) | 0.023 | 48.384 | * | | | |
| **SSS** | Sequence | 0.054 | 4 | 0.014 | 17.122 | * | **0.0217** |
| | Error (sequence) | 0.029 | 36 | 0.001 | | | |
| | Method | 2.147 | 1.819 | 1.180 | 140.624 | * | **0.8626** |
| | Error (method) | 0.137 | 16.371 | 0.008 | | | |
| | Sequence * method | 0.078 | 4.795 | 0.016 | 16.005 | * | **0.0313** |
| | Error (seq * method) | 0.044 | 43.151 | 0.001 | | | |
| **StrS** | Sequence | 0.030 | 4 | 0.007 | 4.625 | 0.004 | **0.0110** |
| | Error (sequence) | 0.058 | 36 | 0.002 | | | |
| | Method | 2.275 | 2.268 | 1.003 | 100.511 | * | **0.8355** |
| | Error (method) | 0.204 | 20.415 | 0.010 | | | |
| | Sequence * method | 0.081 | 3.744 | 0.022 | 9.612 | * | **0.0297** |
| | Error (seq * method) | 0.075 | 33.694 | 0.002 | | | |
| **TraS** | Sequence | 0.040 | 4 | 0.010 | 5.250 | 0.002 | **0.0286** |
| | Error (sequence) | 0.069 | 36 | 0.002 | | | |
| | Method | 1.071 | 5 | 0.214 | 82.588 | * | **0.7645** |
| | Error (method) | 0.117 | 45 | 0.003 | | | |
| | Sequence * method | 0.041 | 4.664 | 0.009 | 5.901 | * | **0.0293** |
| | Error (seq * method) | 0.063 | 41.979 | 0.001 | | | |
| **ICVs** | Sequence | 0.041 | 1.948 | 0.021 | 14.826 | * | **0.0212** |
| | Error (sequence) | 0.025 | 17.534 | 0.001 | | | |
| | Method | 1.764 | 1.401 | 1.259 | 237.110 | * | **0.9144** |
| | Error (method) | 0.067 | 12.613 | 0.005 | | | |
| | Sequence * method | 0.022 | 4.469 | 0.005 | 20.412 | * | **0.0114** |
| | Error (seq * method) | 0.010 | 40.218 | * | | | |

**Supplementary Table S8: Statistical tests of within-subject effects and effect sizes from two-way repeated measures ANOVA analyses.** The ANOVA was run for the whole-brain MVF segmentation and for the four semi-automated single-vein segmentations. Depending on the results of Mauchly's Test of Sphericity, the values are provided either with sphericity assumed or Greenhouse-Geisser corrected. P-values < 0.001 are represented by *. Seq: sequence.

| **p-value** | | iLSQR -STAR | iLSQR - MEDI | iLSQR - TGV | iLSQR - dTIK | iLSQR - iTIK | STAR - MEDI | STAR - TGV | STAR - dTIK | STAR - iTIK | MEDI - TGV | MEDI - dTIK | MEDI - iTIK | TGV - dTIK | TGV - iTIK | dTIK - iTIK |
|---|---|---|---|---|---|---|---|---|---|---|---|---|---|---|---|---|
| **Whole-brain MVF segm.** | Full-FC | 0.928 | * | * | * | * | * | * | * | * | 0.392 | * | 0.171 | * | 0.095 | * |
| | TE1-FC | 0.193 | * | 0.001 | * | * | * | * | * | * | 0.022 | * | 0.232 | * | 0.003 | * |
| | TE1-FC-CS | **0.047** | * | * | * | * | * | * | * | * | 0.199 | * | 0.07 | * | 0.012 | 0.001 |
| | No-FC | 0.004 | * | * | * | * | * | * | * | * | 0.012 | * | 0.334 | * | 0.009 | * |
| | No-FC-7ech | 0.103 | * | 0.015 | * | * | 0.001 | 0.12 | * | * | 0.002 | * | 0.001 | * | * | * |
| **SSS** | Full-FC | * | * | 0.006 | * | * | * | * | * | * | * | 0.001 | 0.004 | * | * | * |
| | TE1-FC | * | * | 0.176 | * | * | * | 0.004 | * | * | * | * | 0.007 | * | * | * |
| | TE1-FC-CS | * | * | 0.142 | * | * | * | 0.005 | * | * | * | * | 0.002 | * | 0.001 | * |
| | No-FC | * | * | 0.299 | * | * | * | 0.01 | * | * | * | * | 0.001 | * | * | * |
| | No-FC-7ech | 0.002 | * | 0.069 | * | * | * | 0.002 | * | * | * | * | 0.005 | * | * | * |
| **StrS** | Full-FC | * | * | 0.017 | * | * | * | * | * | * | * | 0.001 | 0.132 | * | 0.013 | * |
| | TE1-FC | * | * | 0.001 | * | * | * | * | * | * | * | 0.001 | 0.185 | * | 0.048 | * |
| | TE1-FC-CS | * | * | 0.013 | * | 0.002 | * | * | * | * | * | 0.002 | 0.036 | * | 0.526 | * |
| | No-FC | * | * | * | * | * | * | * | * | * | 0.008 | * | 0.096 | * | 0.842 | * |
| | No-FC-7ech | * | * | * | * | * | * | * | * | * | * | * | 0.038 | * | 0.292 | * |
| **TraS** | Full-FC | * | * | * | * | * | * | * | * | * | 0.159 | 0.001 | 0.524 | 0.001 | 0.071 | 0.001 |
| | TE1-FC | * | * | 0.001 | * | * | * | * | * | * | 0.011 | 0.108 | 0.861 | 0.004 | **0.041** | 0.032 |
| | TE1-FC-CS | * | * | 0.001 | * | * | * | * | * | * | **0.043** | 0.057 | 0.694 | 0.004 | 0.069 | 0.079 |
| | No-FC | 0.015 | * | 0.003 | * | * | * | 0.002 | * | * | 0.015 | 0.062 | 0.433 | 0.003 | 0.111 | 0.004 |
| | No-FC-7ech | 0.02 | * | 0.001 | * | * | * | 0.001 | * | * | 0.001 | 0.001 | 0.263 | * | * | 0.011 |
| **ICVs** | Full-FC | * | * | * | * | * | * | * | * | * | 0.001 | * | * | * | * | * |
| | TE1-FC | * | * | * | * | * | * | * | * | * | * | * | * | * | * | * |
| | TE1-FC-CS | * | * | * | * | * | * | * | * | * | * | * | 0.004 | * | * | * |
| | No-FC | * | * | * | * | * | * | * | * | * | 0.002 | * | 0.001 | * | * | * |
| | No-FC-7ech | * | * | * | * | * | * | * | * | * | * | * | 0.003 | * | * | * |

**Supplementary Table S9: P-values from paired samples t-tests investigating (statistically significant) differences between QSM reconstruction methods for different acquisition sequences.** The Benjamini-Hochberg procedure was used to control for the false discovery rate. The p-values < 0.05 that lost significance after the Benjamini-Hochberg procedure are marked in bold. The pairs of methods that remained significantly different from each other after controlling for the false discovery rate are highlighted in blue. P-values < 0.001 are represented by *.

| <u>p-value</u> | | Full-FC - TE1-FC | Full-FC - TE1-FC-CS | Full-FC - No-FC | Full-FC - No-FC-7ech | TE1-FC - TE1-FC-CS | TE1-FC - No-FC | TE1-FC - No-FC-7ech | TE1-FC-CS - No-FC | TE1-FC-CS No-FC-7ech | No-FC - No-FC-7ech |
|---|---|---|---|---|---|---|---|---|---|---|---|
| **Whole-brain MVF segm.** | iLSQR | 0.692 | 0.813 | 0.555 | 0.814 | 0.441 | 0.716 | 0.561 | 0.312 | 0.955 | 0.385 |
| | STAR | 0.279 | 0.077 | 0.073 | 0.061 | 0.508 | 0.622 | **0.05** | 0.694 | * | **0.003** |
| | MEDI | 0.969 | 0.446 | 0.274 | 0.648 | 0.631 | 0.136 | 0.705 | 0.146 | 0.755 | 0.185 |
| | TGV | 0.279 | 0.286 | 0.689 | 0.11 | 0.79 | 0.421 | 0.191 | 0.522 | 0.131 | 0.082 |
| | dTIK | 0.178 | 0.257 | 0.505 | **0.002** | 0.484 | 0.318 | **0.001** | 0.761 | **0.001** | * |
| | iTIK | 0.715 | 0.944 | 0.703 | **0.005** | 0.783 | 0.411 | * | 0.625 | **0.002** | **0.002** |
| **SSS** | iLSQR | 0.311 | 0.155 | 0.427 | 0.383 | 0.466 | 0.938 | 0.837 | 0.717 | 0.397 | 0.805 |
| | STAR | **0.003** | **0.004** | 0.473 | **0.015** | 0.397 | 0.228 | 0.651 | 0.168 | 0.45 | 0.184 |
| | MEDI | 0.117 | 0.733 | 0.351 | **0.005** | **0.006** | **0.019** | * | 0.336 | * | **0.001** |
| | TGV | 0.093 | 0.288 | 0.089 | 0.109 | 0.361 | 0.904 | 0.319 | 0.41 | 0.692 | 0.356 |
| | dTIK | 0.263 | **0.014** | **0.009** | * | **0.007** | **0.004** | * | 0.11 | * | * |
| | iTIK | 0.096 | 0.857 | 0.691 | **0.01** | 0.143 | **0.037** | * | 0.91 | * | **0.001** |
| **StrS** | iLSQR | 0.717 | **0.008** | **0.001** | **0.102** | 0.08 | **0.001** | **0.029** | * | **0.012** | **0.042** |
| | STAR | 0.613 | **0.025** | **0.001** | **0.016** | 0.386 | * | **0.003** | **0.001** | **0.008** | **0.031** |
| | MEDI | **0.031** | 0.559 | 0.378 | 0.243 | **0.024** | 0.267 | **0.006** | 0.621 | 0.136 | **0.023** |
| | TGV | 0.115 | 0.581 | 0.575 | 0.54 | 0.117 | **0.047** | **0.021** | 0.087 | 0.138 | 0.955 |
| | dTIK | 0.058 | 0.943 | 0.882 | **0.017** | **0.02** | 0.079 | **0.001** | 0.77 | **0.009** | **0.008** |
| | iTIK | **0.008** | **0.005** | **0.003** | 0.317 | 0.854 | 1 | **0.003** | 0.845 | **0.002** | **0.002** |
| **TraS** | iLSQR | 0.249 | 0.787 | 0.251 | 0.574 | **0.03** | 0.754 | 0.548 | 0.058 | 0.057 | 0.364 |
| | STAR | 0.766 | 0.587 | 0.962 | 0.153 | 0.146 | 0.768 | **0.014** | 0.745 | 0.149 | 0.226 |
| | MEDI | 0.679 | 0.901 | 0.594 | **0.027** | 0.126 | 0.291 | **0.004** | 0.657 | **0.031** | 0.061 |
| | TGV | **0.045** | 0.248 | 0.109 | 0.068 | 0.16 | 0.741 | 0.522 | 0.403 | 0.192 | 0.948 |
| | dTIK | **0.036** | 0.112 | 0.375 | **0.036** | 0.22 | 0.102 | **0.001** | 0.26 | **0.001** | **0.003** |
| | iTIK | 0.183 | 1 | 0.382 | **0.009** | 0.104 | 0.703 | * | 0.073 | **0.004** | **0.001** |
| **ICVs** | iLSQR | 0.448 | **0.029** | 0.364 | **0.005** | 0.134 | 4.97E-01 | **0.001** | 0.225 | 0.722 | 0.051 |
| | STAR | 0.114 | **0.004** | 0.114 | 0.343 | **0.032** | **0.002** | 0.839 | **0.001** | 0.107 | **0.002** |
| | MEDI | 0.65 | **0.002** | **0.023** | * | **0.021** | **0.035** | * | 0.138 | **0.01** | * |
| | TGV | 0.57 | **0.019** | **0.011** | **0.002** | **0.034** | **0.008** | * | 0.985 | 0.355 | 0.17 |
| | dTIK | 0.818 | **0.004** | 0.168 | * | **0.032** | 0.155 | * | 0.096 | **0.012** | * |
| | iTIK | 0.142 | 0.704 | 0.94 | * | 0.291 | 0.228 | * | 0.709 | **0.006** | * |

**Supplementary Table S10: P-values from paired samples t-tests investigating (statistically significant) differences between acquisition sequences for different QSM reconstruction methods.** All p-values < 0.05 are marked in bold. The pairs of sequences that remained significantly different from each other after controlling for the false discovery rate using the Benjamini-Hochberg procedure are highlighted in blue. P-values < 0.001 are represented by *.

| **Maximal average differences in $\bar{\chi}$-venous** | **MVF** |
|---|---|
| $\Delta\bar{\chi}_{sequence}$ [ppm] | 0.009 |
| $\Delta\bar{\chi}_{method}$ [ppm] | 0.112 |
| $\Delta\bar{\chi}_{method}$ / $\Delta\bar{\chi}_{sequence}$ | 12.44 |

**Supplementary Figure S11: Maximal differences between the method-mean $\bar{\chi}_{method}$ and sequence-mean $\bar{\chi}_{sequence}$ venous susceptibility values and their quotient within the MVF-based automated segmentation from data of the pilot study.** Compared to the main study, the data were acquired in five different subjects on another scanner model from the same vendor. The method-mean values were calculated by averaging over venous susceptibility values acquired with three different sequences but reconstructed with the same QSM method. Accordingly, sequence-mean values were calculated by averaging over venous susceptibility values acquired with the same sequence but reconstructed with six different QSM methods.
